# Distinct stress-dependent signatures of cellular and extracellular tRNA-derived small RNAs (tDRs)

**DOI:** 10.1101/2021.09.03.458085

**Authors:** Guoping Li, Aidan Manning, Alex Bagi, Xinyu Yang, Jonathan Howard, Patricia P. Chan, Thadryan Sweeney, Haobo Li, Brice D. Laurent, Maria I. Kontaridis, Louise C. Laurent, Kendall Van Keuren-Jensen, Sary F. Aranki, Jochen Muehlschlegel, Robert Kitchen, Todd M. Lowe, Saumya Das

**Affiliations:** Cardiovascular Research Center, Massachusetts General Hospital and Harvard Medical School, Boston, MA, USA; Department of Biomolecular Engineering, Baskin School of Engineering, University of California, Santa Cruz, Santa Cruz, CA, USA; Fangshan Hospital of Beijing, University of Traditional Chinese Medicine, Beijing, China; Department of Biomedical Research and Translational Medicine, Masonic Medical Research Institute, Utica, NY, USA; Department of Biological Chemistry and Molecular Pharmacology, Harvard Medical School, Boston, MA, USA; Department of Medicine, Division of Cardiology, Beth Israel Deaconess Medical Center, Harvard Medical School, Boston, MA, USA; Department of Obstetrics, Gynecology, and Reproductive Sciences, University of California, San Diego, La Jolla, CA, USA; Division of Neurogenomics, The Translational Genomics Research Institute, Phoenix, AZ, USA; Division of Cardiac Surgery, Department of Surgery, Brigham and Women’s Hospital, Harvard Medical School. Boston, MA 02115, USA; Department of Anesthesiology, Perioperative and Pain Medicine, Brigham and Women’s Hospital and Harvard Medical School, Boston, MA, USA

## Abstract

The cellular response to stress is an important determinant of disease pathogenesis. Uncovering the molecular fingerprints of distinct stress responses may yield novel biomarkers for different diseases, and potentially identify key signaling pathways important for disease progression. tRNAs and tRNA-derived small RNAs (tDRs) comprise one of the most abundant RNA species in cells and have been associated with cellular stress responses. The presence of RNA modifications on tDRs has been an obstacle for accurately identifying tDRs with conventional small RNA sequencing. Here, we use AlkB-facilitated methylation sequencing (ARM-seq) to uncover a comprehensive landscape of cellular and extracellular tDR expression in a variety of human and rat cells during common stress responses, including nutritional deprivation, hypoxia, and oxidative stress. We found that extracellular tDRs have a distinct fragmentation signature with a predominant length of 31-33 nts and a highly specific termination position when compared with intracellular tDRs. Importantly, we found these signatures are better discriminators of different cellular stress responses compared to extracellular miRNAs. Distinct extracellular tDR signatures for each profiled stressor are elucidated in four different types of cells. This distinct extracellular tDR fragmentation pattern is also noted in plasma extracellular RNAs from patients on cardiopulmonary bypass. The observed overlap of these patient tDR signatures with the signatures of nutritional deprivation and oxidative stress in our cellular models provides preliminary *in vivo* corroboration of our findings and demonstrates the potential to establish novel extracellular tDR biomarkers in human disease models.

## INTRODUCTION

Cells, including unicellular organisms, have evolved sophisticated sensing mechanisms and signal transduction systems to respond to changes in environmental conditions (stress) to ensure the cell survival or alternatively elimination if the cell is unable to cope with the stress^1, 2^. Examples of cellular stress responses include: DNA repair mechanisms triggered by DNA damage during ionizing radiation^3^; the unfolded protein response following heat shock or exposure to chemical toxins^4, 5^; activation of autophagy in response to nutritional deprivation^6^; induction of mitophagy to eliminate damaged mitochondria following hypoxic stress^7^; and adaptive responses to oxidative stress^8, 9^. Increasing evidence has demonstrated that biological processes associated with stress responses play pivotal roles in normal development^10, 11^ and homeostasis^12^, and failure of adaptive stress response can lead to the onset or progression of various diseases^13^. These cellular adaptations to stress involve a complex reorganization of the cellular gene expression program at the level of mRNA biogenesis^1^, which is influenced by the dynamic regulation of non-coding RNA (ncRNA) transcriptome.

As one of the most abundant RNA species in cells, the canonical function of transfer RNAs (tRNAs) in decoding the genetic code in protein translation is well established^14^. More recently, it has been shown that full-length tRNA molecules are processed into smaller regulatory fragments, variously termed tRNA fragments and tRNA halves, or tRNA-derived small RNAs (tDRs), by stress-activated ribonucleases in a regimented manner^15, 16^. tDRs can be grouped into numerous categories. Two of the most prominent types are tRNA halves that are 30-50 nucleotides (nts) long generated by specific cleavage in or close to the anticodon region and tRNA-derived fragments that are usually 12-30 nts in length derived from cleavages of mature or premature tRNAs at various positions^17^. tDRs have been shown to play versatile roles in various biological processes, including gene silencing, RNA stability, protein translation, RNA-binding protein sequestration, epigenetic regulation, and ncRNA regulation^18^. However, most of the existing studies exploring tDR biogenesis and regulation have used hybridization-based methods or conventional RNA sequencing techniques which fail to capture a major portion of the complex tDR pool. Specifically, the presence of tRNA base modifications can hinder the reverse transcription step during small RNA library generation, leading to inaccuracies in both quantification of tDRs and base call accuracy, especially at the ends of sequences^19^. Furthermore, the lack of a widely used, uniform system of nomenclature, coupled with varied computational approaches to tDR read mapping, has led to difficulty in defining reproduceable tDR signatures that can be easily compared between studies^20, 21^. Advances in RNA sequencing methodologies, especially those customized for the RNA modifications commonly seen in tRNAs and tDRs, have helped to overcome the technical challenge of acquiring high quality data. These techniques, including AlkB-facilitated methylation sequencing (ARM-seq) and demethylase-thermostable group II intron RT tRNA sequencing (DM-tRNA-seq), incorporate pretreatment of the input RNA samples with the AlkB enzyme to remove the modifications such as m^1^A, m^1^G and m^3^C on tRNAs and tDRs in order to minimize stalling of reverse transcriptase at modified sites^19, 22^.

As mediators of intercellular communication, extracellular RNAs (exRNAs) have emerged as promising biomarkers for the diagnosis and prognosis of various diseases from minimally invasive liquid biopsies^23, 24^. In addition to high abundance in cells, tDRs also comprise a significantly proportion of the extracellular RNAome^25, 26^. This has been documented in multiple human biofluids, including urine, serum, plasma, saliva, and cerebrospinal fluid, and in cell-conditioned medium^24, 27, 28^. Recent studies suggest a large proportion of extracellular tDRs in plasma or serum, notably the 5’-tRNA halves of certain tRNA isodecoders are associated with ribonucleoproteins^24^. There is also compelling evidence of tDRs associated with EVs in other biofluids and cell culture medium^29, 30^. While the presence of full-length tRNAs within EVs and the site of tDR biogenesis remain topics under current investigation^31, 32^, there does appear to be a correlation between intracellular abundance of tRNAs and their presence in EVs^33^. Recently, several studies have indicated that circulating tDRs could serve as potential biomarkers for cancer diagnosis, prognosis after oncological therapies, monitoring cancer progression, liver fibrosis diagnosis, and distinguishing between subtypes of acute stroke^24, 34^. Because conventional methods for detecting tDRs may underestimate the diversity and abundance of extracellular tDRs, building the stress-specific extracellular tDR signatures using specialized tDR sequencing techniques holds promise to provide new markers of cellular processes associated with disease pathogenesis.

Here, we systematically profile matched cellular and extracellular tDRs expression using ARM-seq in a variety of human and rat cells under three common stressors, including nutritional deprivation, hypoxia, and oxidative stress. We describe the unique fragmentation pattern of extracellular tDRs and the improved discrimination among different cellular stress responses using the extracellular tDR signatures. In preliminary studies, this distinct extracellular tDR fragmentation pattern and stress-specific extracellular tDR signature was observed in plasma extracellular RNAs from patients on cardiopulmonary bypass (CPB), which is a clinical condition involving acute metabolic and oxidative stress. These findings provide a comprehensive landscape of the dynamics of cellular and extracellular tDRs expression induced by common stressors and demonstrate the remarkable potential of extracellular tDR signatures as biomarkers for human diseases.

## RESULTS

### The establishment of *in vitro* stress response models for tDR expression profiling

To systematically profile the molecular signatures of tDRs under different stress conditions, we focused on three common perturbations: nutritional deprivation (glucose and serum deprivation, GSD), hypoxia, and oxidative stress. To enhance the rigor of the study and determine consistent signatures across different cell types, these three perturbations were induced in four different *in vitro* cell culture systems individually, including human embryonic kidney cells – HEK293 (HEK), human choriocarcinoma cells – BeWo, primary neonatal rat ventricular cardiomyocytes (CM), and primary neonatal rat ventricular cardiac fibroblasts (CF) (Fig. 1A).

**Figure 1.**
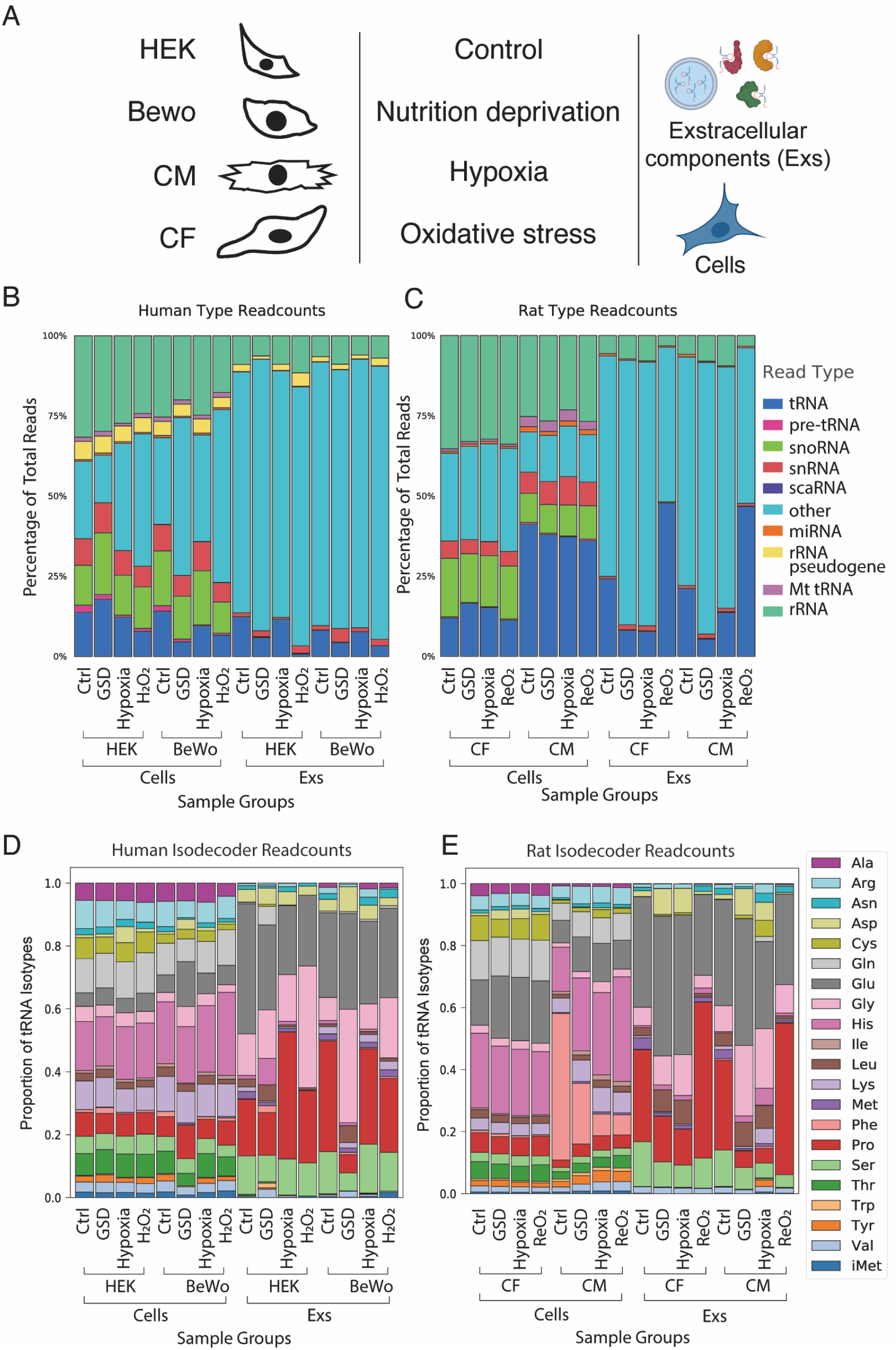
ARM-seq reveals robust information about the cellular and extracellular tDR expression profiles during stress response. **A**. Schematic representation of the samples collected for ARM-seq. **B** and **C**. ARM-seq detects decent amount of tDRs reads from both cells and Exs in human (B) and rat (C) samples; “others” includes mRNA, lincRNAs and all of the other RNA species. **D** and **E**. Extracellular tDRs have isodecoder preferences from tRNA-Glu, tRNA-Gly and tRNA-Pro that are distinct from intracellular tDRs in both human (D) and rat (E) samples.

HEK and BeWo cells were exposed to each stressor for 24 hours. Along with cellular RNAs and proteins, extracellular RNAs were also isolated from conditional cell culture medium using ExoRNeasy kit. Cellular stress responses were confirmed by western blot and qPCR. As expected, GSD significantly activated autophagy and the expression of Ddit3 and GRP78^35^ (Fig. S1A-S1C). Hypoxia stabilized the HIF1a protein and transactivated the expression of Vegf, GluT1, and Ddit4^36^ (Fig. S1A, S1D, and S1E). Oxidative stress, induced by exposure to hydrogen peroxide (H_2_O_2_), significantly induced DNA damage and activated the expression of Ddit3 and Ctgf^37^ (Fig. S1A, S1F, and S1G). To mimic the cardiac ischemia/reperfusion injury, CM and CF were exposed to GSD and hypoxia (0.2% O_2_ in GSD condition) for 5 hours and reoxygenated for an additional 24 hours^38^. Strikingly, there was a rapid response upon GSD or hypoxia treatment, and reoxygenation (ReO_2_) eliminated most of the hypoxia-responsive genes in both CMs and CFs (Fig. S1G and S1H). These data provide strong support for our established *in vitro* stress response models.

### ARM-seq greatly increases the abundance and diversity of tDRs detected

Previously, we developed the ARM-seq platform which improves the accuracy of detection of methylated tDRs^19^. To confirm the improvement of ARM-seq for the detection of tDRs, small RNAs isolated from HEK cells under different stressors were treated with or without purified recombinant His-AlkB and then subjected to deep sequencing after small RNA library preparation. Mapping, annotation, and analysis confirmed that AlkB treatment dramatically increases the proportion of small RNA sequencing reads from tRNA genes (Fig. S2A). Notably, reads after AlkB treatment extend through the ubiquitously modified m^1^A at position 58 of most tRNAs (Fig. S2B). Most importantly, ARM-seq provided an abundance of high-resolution information about the dynamic regulation of tDRs during the stress response, including nutrient deprivation-increased middle fragments of tRNA-Phe-GAA-2/3 and hypoxia-induced 5’tDRs of tRNA-Asp-GTC-2 (Fig. S2C), which were almost undetected by standard small RNA-seq. As a result, we conclude that ARM-seq performed using our purified recombinant His-AlkB works efficiently and facilitates accurate and robust detection of tDRs.

### Overview of intracellular and extracellular tDR expression during the stress response

Next, a total of 96 bar-coded small RNA libraries, including 3 independent replicates for each of the 32 conditions (Fig. 1A), were prepared and sequenced from our *in vitro* stress response models using ARM-seq. Approximate 95.65% of the reads from human samples mapped to the human genome and 95.57% of the reads from rat samples were mapped to the rat genome. Abundant tRNA reads were acquired from both cells and the extracellular components (Exs), including extracellular vesicles (EVs) and ribonucleoproteins, derived from these cells (Fig. 1B and 1C). Interestingly, almost 35% of sequencing reads were aligned to tRNA genes in CMs while this proportion was about 15% in other cell types, which indicates CMs may have higher relative tDR expression (Fig. 1B and 1C). In addition, reoxygenation dramatically increased the proportion of tRNA reads in both CF and CM-derived Exs from ∼25% to ∼50% (Fig. 1C), which suggests an important regulation of extracellular tDRs expression in the heart cells after ischemia/reperfusion. The cellular tRNA isodecoders had similar profiles among different cell types, with the exception of the CMs, which had a higher proportion of tRNA-Phe (Fig. 1D and 1E). The extracellular tRNA isodecoders were also quite similar among the Exs derived from different cell types but had higher proportions of tRNA-Glu and tRNA-Pro and lower proportions of tRNA-Arg, tRNA-Gln, and tRNA-His, when compared with cellular tRNA isodecoders (Fig. 1D and 1E).

### Intracellular and extracellular tDRs have distinct fragmentation signatures

Increasing evidence has shown that tDRs are a key component of extracellular RNAs and account for the majority of mapped reads in most biofluids tested^27, 39^. However, the fragmentation profiles of intracellular and extracellular tDRs have not been systematically studied. Here, we first analyzed the size of tDRs among the 96 samples. Strikingly, the extracellular tDRs were predominantly 31-33 nts in length across the species, cell types, and stressors, while the intracellular tDRs had a wide range of length distribution (Fig. 2A-2D and Fig. S3A-S3L). This predominance for a specific length appeared to be specific to tDRs, and was not seen in other small extracellular RNA types (Fig. S4A-S4P). 31-33 nts tDRs are usually considered as tRNA-derived halves^40^. To further confirm the subtypes of these specific extracellular tDRs, the read coverage across each position of tRNAs were analyzed based on the reads of each nucleotide. We found that around 70% of the extracellular tDRs are aligned to the 5’ halves of tRNAs with about 20% aligned to the 3’ end of tRNAs, whereas the intracellular tDRs are specifically derived from the 3’ end of tRNAs (Fig. 2E-2H and Fig. S5A-S5L). The termination position analysis further confirmed that the extracellular 5’tDRs end at the anti-codon loop which usually generates 5’ tRNA halves (Fig. 2I-2L and Fig. S6A-S6I). Notably, almost all the extracellular tDRs derived from the 3’ end of tRNAs had a trimmed end at position 74 ending with cytosine, while most of the cellular 3’tDRs had an intact CCA end (Fig. 2I-2L and Fig. S6A-S6L), suggesting that extracellular 3’tDRs may have a specific biogenesis mechanism that is distinct from their intracellular counterparts. In summary, these data strongly demonstrated that tDRs have distinct fragmentation signatures between cells and Exs.

**Figure 2.**
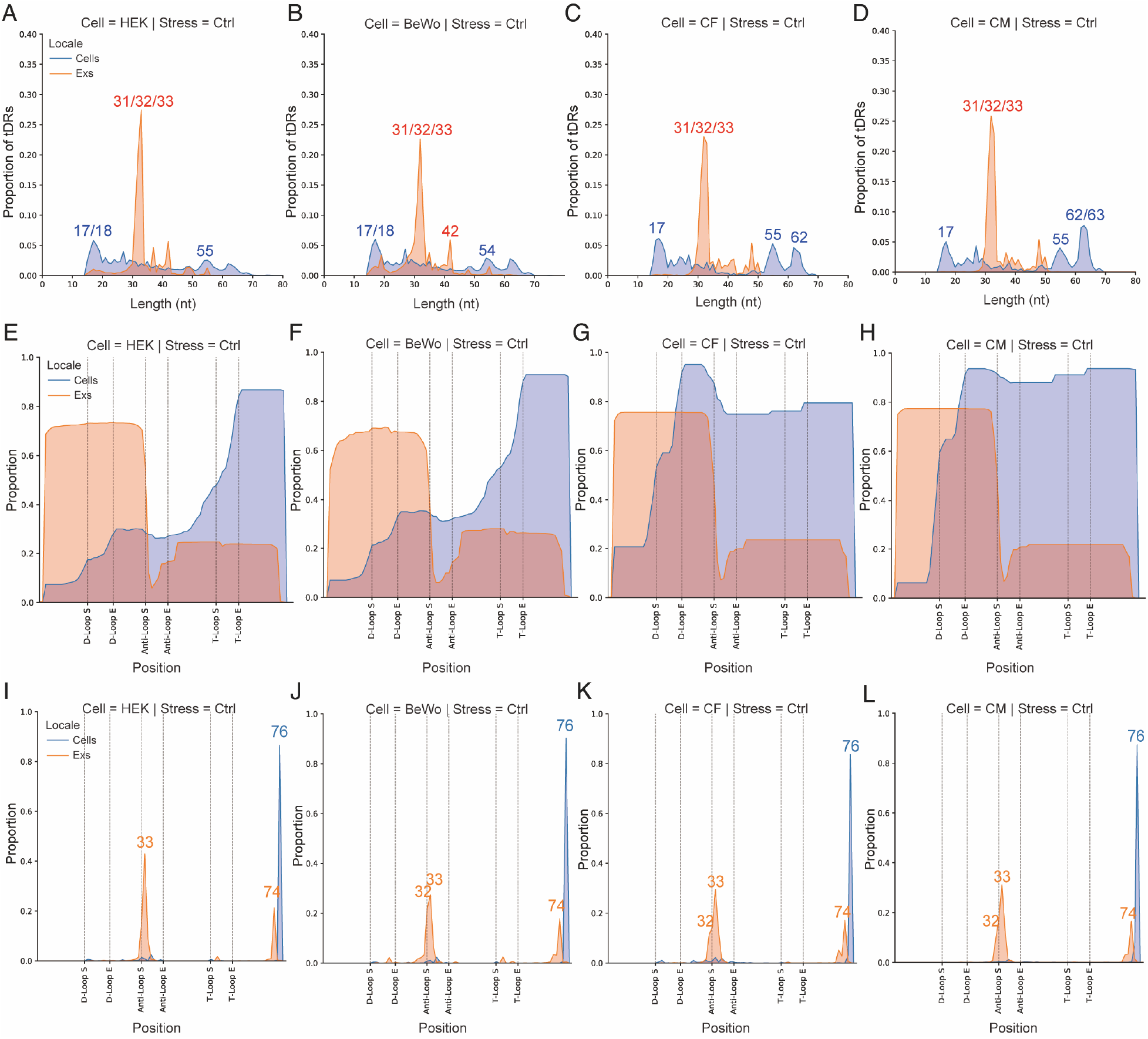
Extracellular tDRs show distinct fragmentation signatures from cellular tDRs. **A-D**. Extracellular tDRs are predominantly 31-33nt in length in all profiled cell types, including HEK (A), BeWo (B), CF (C), and CM (D). **E-H**. Extracellular tDRs are predominantly tRNA halves and derived from both ends of tRNA genes while intracellular tDRs are mainly generated from 3’ end of tRNAs with various lengths in all cell types tested, including HEK (E), BeWo (F), CF (G), and CM (H). **I-L**. Extracellular tDRs end at either position 33 or 74 while most of the intracellular tDRs end at position 76 in all examined cell types, including HEK (I), BeWo (J), CF (K), and CM (L) cells.

### Extracellular tDR expression profiles provide better discrimination between different stress responses compared to miRNAs

Extracellular miRNA expression has been extensively studied to identify biomarkers for the diagnosis and prognosis of diseases^41^, whereas our knowledge of extracellular tDRs is still emerging. To evaluate the capability of tDRs to distinguish between different stress responses, we assessed the differences in the expression levels of different small RNA species, including tDRs and miRNAs, across all profiled samples from each cell type by performing principal component analysis (PCA) clustering. Interestingly, when clustering the 24 samples from BeWo cells based on tDR signatures, cellular samples subjected to different treatments were largely overlapping, whereas different stressor-induced EV tDR profiles were clearly distinguished from each other (Fig. 3A). In contrast, clustering based on extracellular miRNA expression profiles did not separate samples according to exposure to stressors, while the cellular miRNA expression signatures were able to distinguish between them (Fig. 3B). The unique property of extracellular tDR expression profiles to discriminate different stress responses were also demonstrated in HEK (Fig. S7A and S7B), CM (Fig. 3C and 3D), and CF (Fig. S7C and S7D) cells.

**Figure 3.**
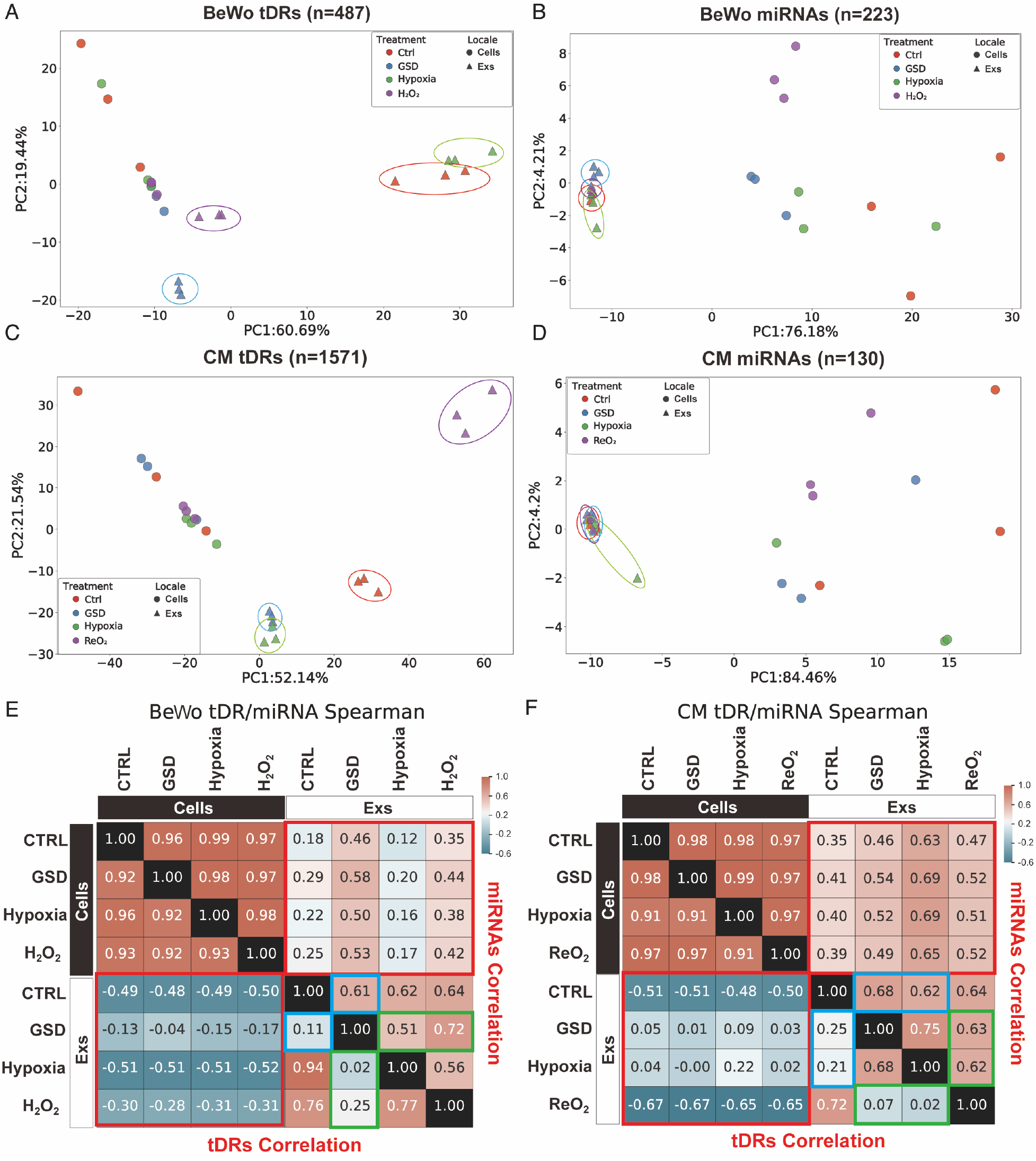
Extracellular tDR expression landscapes provide better discrimination between different stress responses compared to miRNAs. **A** and **B**. PCA analysis of tDR profiles (A) provides better resolution to distinguish EV samples (circled) derived from BeWo cells after different stress treatments than miRNA profiles (B). **C** and **D**. PCA analysis of tDR profiles (C) provides better resolution to distinguish EV samples (circled) derived from CM cells after different stress treatments than miRNA profiles (D). **E.** Heatmaps of correlation coefficients (Spearman) for tDR class (left bottom) shows larger variance among different samples than miRNA class (right top) in BeWo cells. **F.** Heatmaps of correlation coefficients (Spearman) for tDR class (left bottom) shows larger variance among different samples than miRNA class (right top) in CM cells. In E and F, red boxes show the difference between intracellular samples and extracellular samples, blue boxes indicate the difference between each stressor and control group in EV samples, and green boxes present the difference between different stressors in EV samples.

To further support these findings, we performed dimensionality reduction via Uniform Manifold Approximation and Projection (UMAP). As expected, the UMAP for tDR signatures clearly delineated each stressor-exposed extracellular RNA sample from extracellular RNA samples exposed to other stressors in each profiled cell type, while the extracellular miRNA expression profiles had a limited capacity to discriminate among different stress treatments (Fig. S8A-S8H). Finally, similar to the pattern observed for miRNA expression, tDR expression was highly correlated across the 4 different cellular samples, but showed poor correlation between cellular and extracellular samples (Fig. 3E, 3F, S9A, and S9B), indicating significant differences between cellular and extracellular small RNA expression profiles. The tDR expression profiles appeared to be more divergent between the extracellular and cellular compartments and between the different stress responses compared to the corresponding miRNA expression profiles (Fig. 3E, 3F, S9A, and S9B). Taken together, our data strongly suggested that the extracellular tDRs expression signatures are not simply a reflection of the stoichiometry of cellular tDR expression, and provide improved discrimination between different stress responses than miRNA signatures.

### Nutritional deprivation-shaped cellular and extracellular tDR signatures

Stress, especially amino acid starvation, has been shown to induce dynamic expression of tDRs^20^. However, the abundance and diversity of tDRs were previously underestimated due to the technical shortcomings of conventional small RNA-sequencing. To systematically study the regulation of tDRs after nutrition deprivation, we assessed differentially expressed cellular and extracellular tDRs in response to nutrition deprivation in all four cell types. By tracking the expression of each tDR in HEK cells or in Exs with or without GSD treatment, we again noted the distinct expression profiles of cellular and extracellular tDRs (Fig. 4A), confirming our previous conclusion that extracellular tDRs do not simply reflect the stoichiometry of cellular tDRs. Strikingly, extracellular tDRs were far more dynamically regulated after GSD treatment when compared with cellular tDRs (Fig. 4A). In contrast, there were fewer expressed extracellular miRNAs with or without GSD treatment, although the cellular miRNA expression levels were comparable to the level of tDRs (Fig. 4B). These findings were also noted in BeWo (Fig. S10A and S10B), CM (Fig. 4C and 4D), and CF cells (Fig. S10C and S10D). Together, our results demonstrate that the biogenesis or packaging of extracellular tDRs (and to a lesser extent, cellular tDRs) is dynamically regulated by the cellular response to nutrient deprivation and may provide a more sensitive marker of metabolic stress than miRNAs.

**Figure 4.**
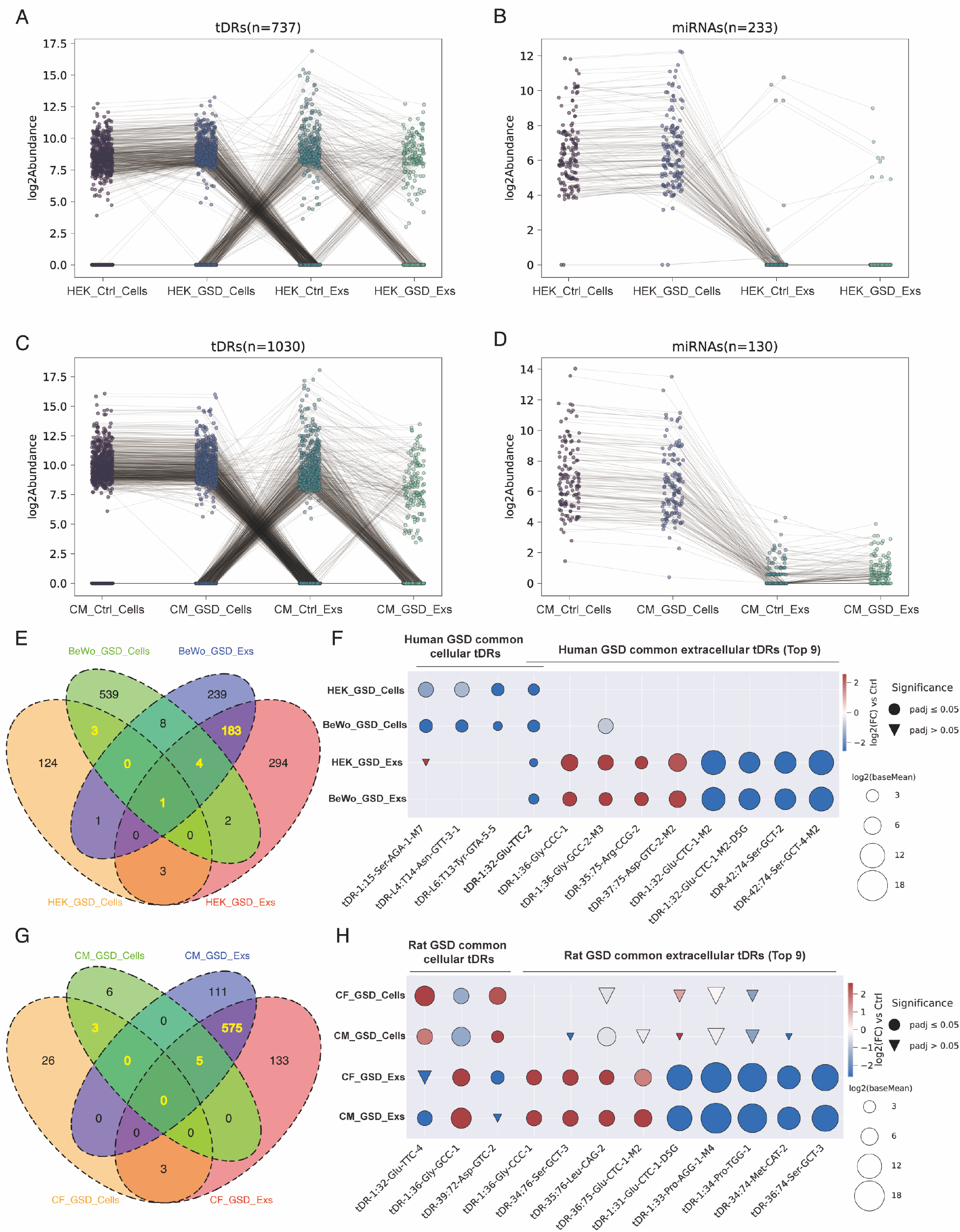
Nutritional deprivation-shaped cellular and extracellular tDR signatures. **A** and **B**. Expression tracing plots reveal more dynamic changes of extracellular tDRs (A) than miRNAs (B) in HEK cells during GSD treatment. **C** and **D**. Expression tracing plots reveal more robust changes of extracellular tDRs (C) than miRNAs (D) in CM cells during GSD treatment. **E.** Venn diagram shows the overlapped and specific tDRs that were regulated by GSD in human cells and Exs. **F.** The most significant cellular and extracellular tDRs that were regulated by GSD in both HEK and BeWo cells. **G.** Venn diagram shows the overlapped and specific tDRs that were altered by GSD in rat cells and Exs. **H.** The most significant cellular and extracellular tDRs that were altered by GSD in both CF and CM cells.

In HEK cells, DESeq analysis discovered 111 upregulated and 21 downregulated cellular tDRs and 120 upregulated and 367 downregulated extracellular tDRs after GSD treatment (Table. S1). In BeWo cells, there were 9 upregulated and 548 downregulated cellular tDRs and 84 upregulated and 352 downregulated extracellular tDRs after GSD treatment (Table. S1). Notably, only 4 differentially expressed cellular tDRs were common to both HEK and BeWo cells with nutritional deprivation (Fig. 4F). As opposed to the cellular tDRs, there were 188 common differentially expressed extracellular tDRs upon GSD treatment (Fig. 4E) of which 27 were upregulated and 161 were downregulated (Table S1). The top 4 significantly upregulated extracellular tDRs associated by both cell types are tDR-1:36-Gly-CCC-1, tDR-1:36-Gly-GCC-2-M3, tDR-35:75-Arg-CCG-2, and tDR-37:75-Asp-GTC-2-M2 while the top 4 significantly downregulated extracellular tDRs are tDR-1:32-Glu-CTC-1-M2, tDR-1:32-Glu-CTC-1-M2-D5G, tDR-42:74-Ser-GCT-2, and tDR-42:74-Ser-GCT-4-M2 (Fig. 4F). tDR-1:32-Glu-TTC-2 was downregulated in both cells and extracellular compartment in response to GSD for both cell types.

In the rodent primary cell cultures cultured in GSD for 5 hours, DESeq2 analysis identified only 32 differentially expressed cellular tDRs in neonatal rat CFs and 14 changed tDRs in CMs, of which 3 were common between the two cell types: tDR-1:32-Glu-TTC-4, tDR-1:36-Gly-GCC-1, and tDR-39:72-Asp-GTC-2 (Fig. 4G, Fig. 4H and Table S2). In contrast, there were 716 and 691 differentially expressed extracellular tDRs for CFs and CMs after GSD, respectively, of which around 10% of these were upregulated (Fig. 4G and Table S2). Of the 580 extracellular tDRs that were differentially expressed in both cell types, 44 were upregulated (Table S2). The top 4 significantly upregulated and top 4 significantly downregulated extracellular tDRs associated by both cell types are enumerated in Figure 4H. Interestingly, we found a number of conserved extracellular tDRs among rat and human species that were commonly regulated by nutritional deprivation across all four cell types, including the upregulated tDR-1:36-Gly-CCC-1, tDR-1:35-Gly-CCC-1, tDR-1:36-Asp-GTC-2, tDR-1:36-Glu-CTC-1-M2-C35U, and tDR-3:36-Gly-GCC-1, and approximately 85 downregulated tDRs, such as tDR-1:16-Leu-AAG-1-M4, tDR-1:29-Pro-AGG-1-M6, tDR-1:30-Glu-CTC-1-M2, tDR-1:31-Gly-GCC-1, tDR-1:32-His-GTG-1, tDR-37:74-Asp-GTC-2-M2, and tDR-39:74-Glu-CTC-1-M2 (Tables S1 and S2; listed as human annotations). These data suggest that there may be a ‘universal’ extracellular tDR signature of nutritional deprivation.

### Hypoxia-shaped cellular and extracellular tDR signatures

The cellular response to hypoxia plays a key role in the pathogenesis of many diseases, including myocardial ischemia, metabolic disorders, chronic heart and kidney diseases, and reproductive diseases^42^. To comprehensively profile the tDR signature of the cellular response to hypoxia, we compared the expression levels of cellular and extracellular tDRs between normoxic and hypoxic conditions in the four cell types. Consistent with our findings for nutritional deprivation, RNA expression tracking plots showed more dramatic changes in the expression of extracellular tDRs expression than intracellular tDRs expression and more pronounced changes in tDRs than miRNAs, in all profiled cell types upon hypoxia treatment (Fig. S11A-S11H).

Unlike the nutritional deprivation response, alternations in tDRs expression in response to hypoxia were less pronounced, with 26 differentially expressed cellular tDRs and 55 changed extracellular tDRs in BeWo cells, and 151 differentially expressed cellular tDRs and 309 changed extracellular tDRs in HEK cells (Fig. S12A and Table S3). We noted an increase of tDR-1:34-Gly-GCC-1, which was previously reported to be induced by hypoxia in triple-negative breast cancer cells^43^, in hypoxia-treated BeWo cells but not in HEK cells (Table S3). Strikingly, the 13 cellular tDRs that were altered by hypoxia in both HEK and BeWo cells were all downregulated, within which the top 4 tDRs were tDR-T1:T20-Ser-TGA-1-1, tDR-38:T9-Arg-TCT-1-1, tDR-1:32-Pro-AGG-1-M5, and tDR-38:T16-Tyr-GTA-2-1 (Fig. S12B and Table S3). There were 16 hypoxia-regulated extracellular tDRs common to HEK and BeWo cells, among which the top 4 upregulated tDRs were tDR-14:33-Glu-TTC-1-M2, tDR-39:73-Glu-TTC-2, tDR-42:73-Arg-CCT-4, and tDR-7:33-Glu-TTC-1-M2 and the 4 most significantly downregulated tDRs were tDR-1:33-Glu-CTC-1-M2, tDR-1:33-Glu-CTC-1-M2-D5G, tDR-1:33-Gly-GCC-1, and tDR-34:74-Met-CAT-3 (Fig. S12B and Table S3). tDR-1:33-Lys-CTT-2 was downregulated by hypoxia treatment in both cells and Exs from HEK and BeWo (Fig. S12B and Table S3).

The differential expression analysis revealed more pronounced changes in extracellular tDRs in hypoxia-treated rat CF and CM samples, with 724 significantly changed extracellular tDRs associated with CF and 688 differentially expressed tDRs associated with CMs (Fig. S12C and Table S4). 58 out of the 511 extracellular tDRs that were changed in both cell types were upregulated; the top 4 of these tDRs were tDR-1:36-Gly-CCC-1, tDR-21:75-Cys-GCA-1-M2, tDR-35:76-Leu-CAG-2, and tDR-37:76-Glu-CTC-1-M2 (Fig. S12C, Fig. S12D and Table S4). The 4 most significantly common-downregulated extracellular tDRs were tDR-1:33-Pro-AGG-1-M4, tDR-1:34-Pro-TGG-1, tDR-36:74-Ser-GCT-3, and tDR-37:74-Glu-TTC-3-M2 (Fig. S12D and Table S4). In the cells, there were 105 differentially expressed tDRs in CM cells but only 27 tDRs were changed in CF cells after hypoxia treatment (Fig. S12C and S12D). 6 of them overlapped and were upregulated in both CF and CM cells, within which the top 4 tDRs were tDR-1:32-Glu-CTC-1, tDR-1:32-Glu-TTC-4, tDR-1:33-Glu-TTC-4, and tDR-39:72-Asp-GTC-2 (Fig. S12D and Table S4). Interestingly, most of these upregulated cellular tDRs were significantly downregulated in CF and CM Exs after the treatment of hypoxia (Fig. S12D). Notably, we also found some conserved extracellular tDRs among human and rat species that were downregulated by hypoxia in the extracellular compartment for all four cell types, including tDR-1:33-Glu-CTC-1, tDR-1:33-Glu-CTC-1-D5G, tDR-18:33-Glu-CTC-1-M2, tDR-2:33-Glu-CTC-1-C4U-U5G, tDR-34:74-Met-CAT-2, tDR-34:74-Met-CAT-3, tDR-35:72-Leu-CAG-2, and tDR-37:74-Glu-CTC-1-M2 (Table S3 and S4; listed as human annotations). Similar to the response of the cells to nutritional deprivation, these data suggest that the regulation of the biogenesis of extracellular tDRs remains distinct from that of cellular tDRs in all the cell types examined. Furthermore, there appear to be several key ‘common’ extracellular signatures associated with hypoxia in the rodent and human-derived cell lines.

### Oxidative stress-shaped cellular and extracellular tDR signatures

The generation of tDRs through tRNA cleavage has been shown to be a conserved response to oxidative stress in eukaryotes^44^. Hence, we also characterized the oxidative stress-specific tDR signatures in the four cell types to. Interestingly, the expression levels of most of the differentially expressed tDRs, including both cellular and extracellular tDRs, were decreased upon H_2_O_2_ treatment in HEK and BeWo cells; in contrast, oxidative stress induced by ReO_2_ led to a significant increase of extracellular tDRs in both CM and CF cells. Cellular miRNAs or extracellular miRNAs only demonstrated minor changes during oxidative stress response (Fig. S13A-S13H).

The majority of cellular tDRs in HEK and BeWo cells after H_2_O_2_ treatment were downregulated (Fig. S13A, Fig. S13C and Table S5). The expression levels of all 188 cellular tDRs that were significantly changed in both HEK and BeWo cells were reduced by oxidative stress, including tDR-58:76-Cys-GCA-1-M9, tDR-L4:T12-Val-AAC-1-4, tDR-L4:T13-Arg-CCT-4-1, tDR-L4:T14-Asn-GTT-2-8, tDR-L4:T14-Lys-CTT-1-1, and tDR-1:32-Pro-AGG-1-M5, as the top 6 most significant tDRs (Fig. S14A, Fig. S14B and Table S5). In the extracellular samples, all the 423 differentially expressed extracellular tDRs from HEK cells and 298 out of 305 extracellular tDRs from BeWo cells were significantly downregulated during oxidative stress response (Fig. S14A and Table S5). Of the 157-overlapping extracellular tDRs associated with both cell types, the top 6 are shown in Figure S14B (Fig. S14B and Table S5).

As detailed above, to better phenocopy the oxidative stress in models of cardiac ischemia/reperfusion, our model of oxidative stress for CMs and CFs involved exposure to hypoxia (0.2% O_2_ in GSD condition) for 5 hours and ReO_2_ for an additional 24 hours. Unexpectedly, there were no differentially expressed tDRs for CF cells but there were 11 significantly changed tDRs in CM cells after ReO_2_ (Fig. S14C and Table S6). In CMs, the levels of tDR-1:32-Glu-TTC-4, tDR-36:73-Asn-GTT-1, tDR-3:32-Gly-CCC-1-M2, tDR-3:32-Gly-GCC-2-M2, and tDR-21:74-His-GTG-1 were significantly increased, and tDR-1:34-Glu-CTC-1-D5G, tDR-12:76-Glu-TTC-3-M2, tDR-15:76-Glu-TTC-3-M2, tDR-11:76-Glu-TTC-3-M2, tDR-1:26-Glu-TTC-4, and tDR-1:20-Pro-AGG-1-M5 were downregulated after ReO_2_ (Table S6). In contrast to the changes noted with the HEK and BeWo cells, 548 and 561 extracellular tDRs from CM and CF cells were dramatically altered after ReO_2_, and most of them were upregulated (Fig. S14C and Table S6), consistent with the tDR expression tracing plots (Fig. S13E-S13H). 388 overlapping extracellular tDRs were significantly modulated by ReO_2_ in both CM and CF cells and only 16 of them had decreased expression levels (Fig. S14C and Table S4). The top 5 downregulated extracellular tDRs and the top 7 upregulated extracellular tDRs in both cell types after the treatment of ReO_2_ are enumerated in Figure S14D. Of interest, we noticed decreased expression levels of tDR-1:36-Glu-CTC-1 and tDR-1:36-Glu-CTC-1-D5G and the increased levels of tDR-2:31-Glu-CTC-1 and tDR-2:31-Glu-CTC-1-D4G in both CM and CF Exs in response to ReO_2_ (Fig. S14D); the distinct regulation of these tDRs derived from the same parent tDR is suggestive of different modes of biogenesis or differential stability of these fragments. Strikingly, we also uncovered 5 extracellular conserved tDRs among human and rat species that are downregulated by oxidative stress, including tDR-1:33-Val-AAC-1-M6, tDR-1:36-Glu-CTC-1-M2, tDR-21:56-Leu-AAG-2-M3-G14A, tDR-42:74-Arg-CCT-4, and tDR-42:74-Ser-GCT-2 (Tables S5 and S6). Our data strongly suggested that the cellular context is important for interpreting the response to oxidative stress. In the case of all the cell lines, extracellular tDRs were far more altered than cellular tDRs in response to oxidative stress. While the stress models were different between the human and rodent-derived cells, the differences in the directionality of changes in the extracellular tDRs were notable.

### Specific and shared extracellular tDR signatures among three profiled stressors

It is well established that the levels of tDRs in human liquid biopsy can dynamically change under different diseases^24^. However, the knowledge on the basic variables influencing the extracellular tDR levels is still limited. To define the stressor-specific and shared extracellular tDR signatures, Venn diagram of differentially expressed extracellular tDRs identified under three stressors in both HEK and BeWo cells were created (Fig. 5A). Interestingly, more than 60% of GSD-altered extracellular tDRs (118 out of 188) and 50% of hypoxia-shaped extracellular tDRs (8 out of 16) were also regulated by H_2_O_2_ (Fig. 5A). tDR-1:33-Glu-TTC-14-M3 and tDR-34:74-Met-CAT-3 were downregulated in Exs by all three profiled stressors (Fig. 5B). There results indicated that there are shared stress response mechanisms among nutritional deprivation, hypoxia, and oxidative stress. Notably, 27 out of 69 extracellular tDRs that were specifically regulated by GSD in both HEK and BeWo cells were significantly upregulated; the top 6 of these tDRs were tDR-1:36-Gly-CCC-1, tDR-1:36-Gly-GCC-2-M3, tDR-34:75-Ser-GCT-4-M2, tDR-35:75-Arg-CCG-2, tDR-35:75-Leu-CAG-2, and tDR-37:75-Asp-GTC-2-M2 (Fig. 5A and 5B). About 70% of hypoxia specifically altered extracellular tDRs (5 out of 7) were upregulated, including tDR-14:33-Glu-TTC-1-M2, tDR-39:71-Asp-GTC-2-M2, tDR-39:73-Glu-TTC-2, tDR-42:73-Arg-CCT-4, and tDR-7:33-Glu-TTC-1-M2 (Fig. 5A and 5B). All of the 33 H_2_O_2_ specifically shaped extracellular tDRs were downregulated (Fig 5A and 5B).

**Figure 5.**
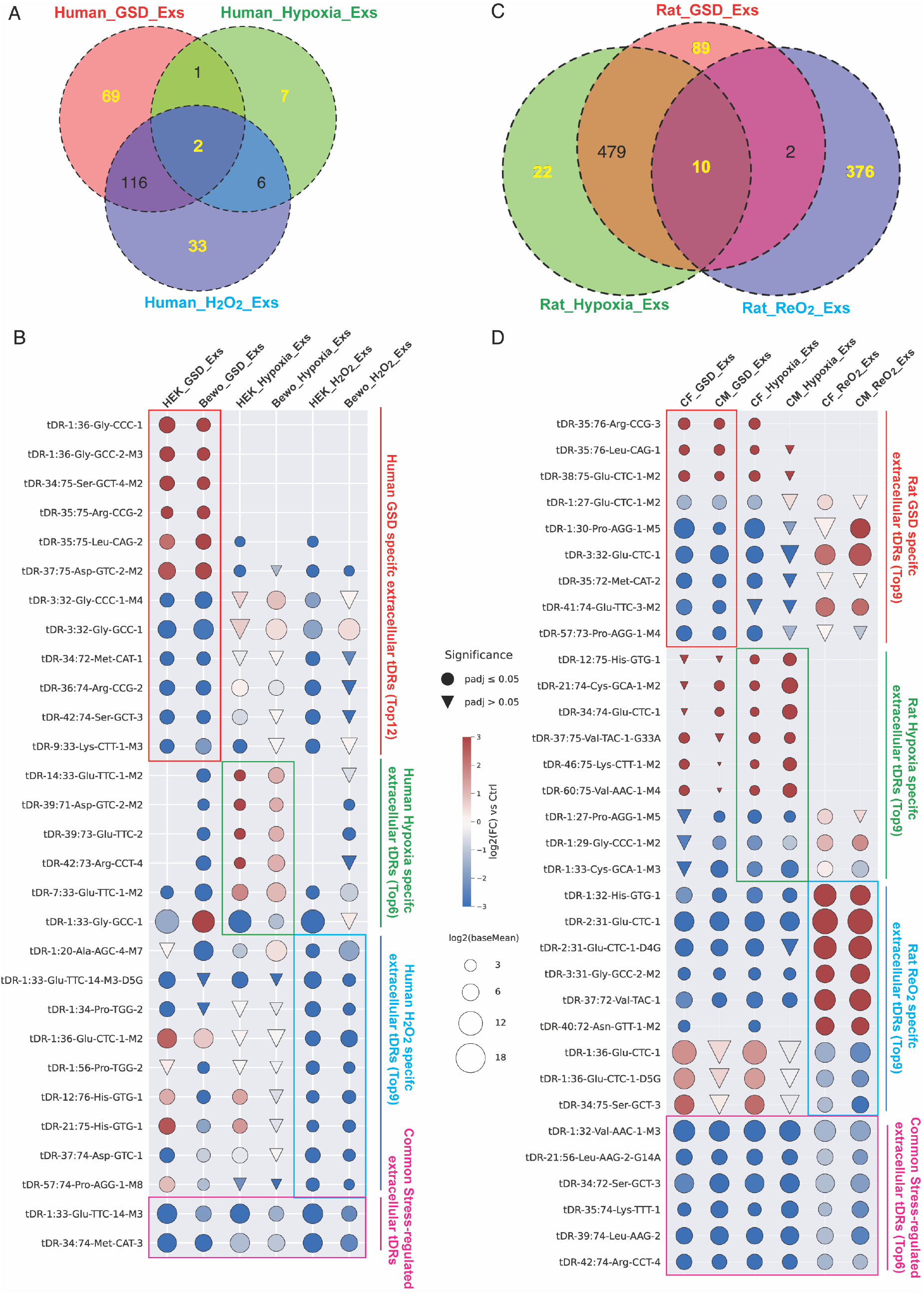
Specific and shared extracellular tDR signatures among three profiled stressors. **A.** Venn diagram shows the specific and shared extracellular tDRs altered by three profiled stressors in HEK and BeWo cell-derived Exs. **B.** The most significantly regulated extracellular tDRs that were specifically for GSD, hypoxia, and H_2_O_2_, and were shared among three stressors in HEK and BeWo cell-derived Exs. **C.** Venn diagram shows the specific and shared extracellular tDRs altered by three profiled stressors in CF and CM cell-derived Exs. **D.** The most significantly regulated extracellular tDRs that were specifically for GSD, hypoxia, and ReO_2_, and were shared among three stressors in CF and CM cell-derived Exs.

The analysis of rodent stress-shaped extracellular tDR signatures identified 10 extracellular tDRs that were significantly downregulated by GSD, hypoxia, and ReO_2_; the top 6 of these tDRs were tDR-1:32-Val-AAC-1-M3, tDR-21:56-Leu-AAG-2-G14A, tDR-34:72-Ser-GCT-3, tDR-35:74-Lys-TTT-1, tDR-39:74-Leu-AAG-2, and tDR-42:74-Arg-CCT-4 (Fig. 5C and 5D). 95% (489 out of 511) of extracellular tDRs that were altered by hypoxia were also significantly regulated by GSD in both CF and CM cells (Fig. 5C), which may due to the hypoxia condition used in primary cardiac cells contains nutritional deprivation as well. Within the 89 GSD specifically regulated extracellular tDRs, only tDR-35:76-Arg-CCG-3, tDR-35:76-Leu-CAG-1, and tDR-38:75-Glu-CTC-1-M2, were upregulated (Fig. 5C and 5D). Interestingly, more than 70% (17 out of 22) of the hypoxia specifically altered extracellular tDRs were upregulated (Fig. 5C and 5D), while approximately 99% (372 out of 376) of the ReO_2_ specifically regulated extracellular tDRs were upregulated (Fig. 5C and 5D). Notably, tDR-1:36-Glu-CTC-1 and tDR-1:36-Glu-CTC-1-D5G were specifically downregulated by oxidative stress in all of the profiled human and rodent cells (Fig. 5B and 5D), which suggest a conservative extracellular tDR signature specifically for oxidative stress.

### Patient plasma tDR signatures reveal distinct stress responses during cardiac surgery with CPB

Cardiac surgery remains one of the most commonly performed major surgeries for the patients with valvular abnormalities or multivessel coronary diseases^45^. During the procedure, the heart is typically arrested and connected to a CPB machine, which provides both perfusion pressure and oxygenation to support the circulation^46^. During the short time period of the CPB, the heart is exposed to metabolic and oxidative stress (especially during reperfusion)^47^. As a pilot ‘test’ case to determine the applicability of our extracellular tDR signatures to human subjects, we collected the plasma samples from human patients at the initiation (Pre-CPB) and about 73 minutes of CPB (Post-CPB). ARM-seq was performed on RNAs isolated from these plasma EV samples. The mapping results showed that around 10% total reads were mapped to tRNA genes in these human plasma samples (Fig. 6A); this detection was far more robust than previously reported for plasma using conventional small RNA-seq in previous studies^27, 48^.

**Figure 6.**
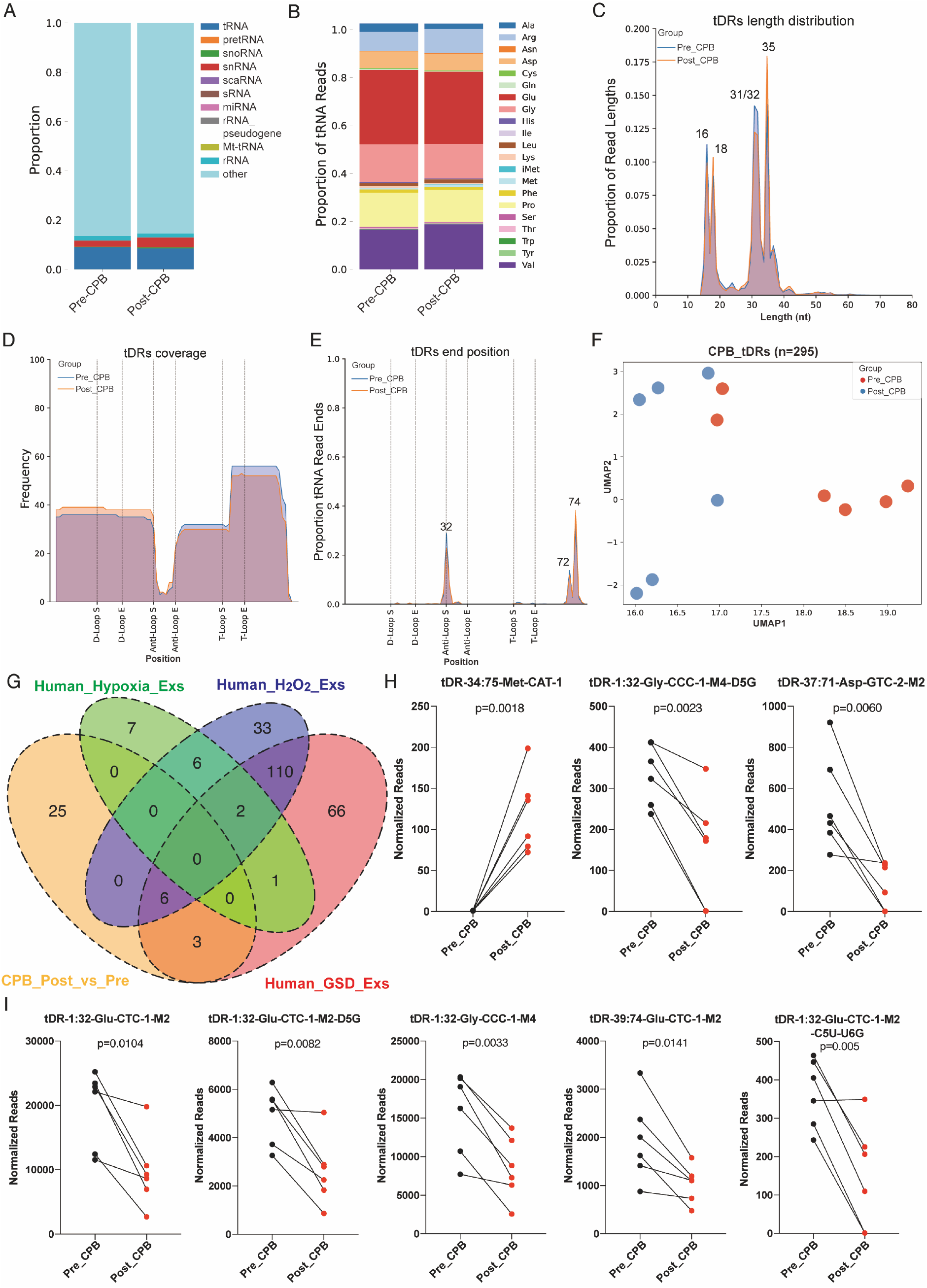
Patient plasma tDR signature reveals distinct stress responses during CPB surgery. **A.** About 10% total reads mapped to tRNA genes in the examined human plasma Exs samples. **B.** A large proportion of plasma tDRs are derived from tRNA-Glu, tRNA-Gly, tRNA-Pro, and tRNA-Val. **C.** Plasma tDRs are mainly 16-18 nts or 31-35 nts in length. **D.** Plasma tDRs are predominantly tRNA halves that derived from both ends of tRNA genes. **E.** Plasma tDRs end at position 32, 72 or 74 of tRNAs. **F.** UMAP projection based on tDR landscapes provides clear resolution to distinguish pre-CPB from post-CPB surgery. **G.** CPB surgery-modulated tDRs are overlapped with nutritional deprivation-shaped and oxidative stress-shaped extracellular tDR signatures but not hypoxia-shaped extracellular tDR signature. **H.** Three CPB surgery-modulated tDRs that were only found in nutritional deprivation-shaped extracellular tDR signatures. Paired two-tailed t-test. **I.** Top 5 CPB surgery-modulated tDRs that were found in both nutritional deprivation-shaped and oxidative stress-shaped extracellular tDR signatures. Paired two-tailed t-test.

The tRNA isodecoder analysis revealed that a large proportion of tDRs in human plasma Exs are derived from tRNA-Glu, tRNA-Gly or tRNA-Pro (Fig. 6B), similar to our findings from cell culture (Fig. 1C). More than 60% of plasma tDRs are 31-35 nts in length and most of them are tRNA halves from either 5’ end or 3’ end of tRNA genes, which end at position 32 (anticodon loop) for the 5’ end, 72 (before the last nucleotide of 3’ end) or 74 (after the first nucleotide of 3’ CCA tail) (Fig. 6C-6E). These plasma tDR fragmentation profiles are consistent with the extracellular tDR fragmentation profile from cell culture (Fig. 2). Although a proportion of tDRs, which had 16-18 nts in length and were generated by dual cleavages at T-loop and position 72, were only found in plasma Exs but not in the Exs from our cell culture systems, our findings still indicates that extracellular tDRs present in the cell culture systems and plasma may share pathways of biogenesis.

UMAP projection analysis based on tDR expression profiles showed modest resolution for distinguishing pre-CPB surgery patients from post-CPB surgery patients based on tDR signatures (Fig. 6F). This suggests that extracellular tDRs may be robust markers of stress response. Next, we selected the tDRs that were significantly changed during CPB and assessed any overlap with our three common-stress-specific extracellular tDRs signatures. 8 out of 17 differentially expressed tDRs after about 73 minutes of CPB surgery were found in the nutritional deprivation-specific extracellular tDR signatures and 5 of them overlapped with both nutritional deprivation-specific and oxidative stress-specific extracellular tDR signatures; none of them were represented in the hypoxia-specific extracellular tDR signatures (Fig. 6G). Notably, among the 3 tDRs that were altered by CPB and nutritional deprivation, tDR-34:75-Met-CAT-1 was only detected in human plasma Exs after about 73 minutes of CPB (Fig. 6H). The tDRs derived from tRNA-Glu-CTC-1, including tDR-1:32-Glu-CTC-1-M2, tDR-1:32-Glu-CTC-1-M2-D5G, tDR-1:32-Glu-CTC-1-M2-C5U-U6G, tDR-2:32-Glu-CTC-1-M2-C4U-U5G, and tDR-39:74-Glu-CTC-1-M2, were significantly downregulated with CPB, in common with exposure to nutritional deprivation or oxidative stress in our cell culture systems (Fig.6I). Overall, these results support prior data suggesting that cells experience oxidative stress and nutritional deprivation during CPB and provides a novel circulating RNA signature for these cellular processes.

## DISCUSSION

The expanding use of RNA sequencing technology has led to an explosion in the discovery of novel RNA species. Notably, considerable new knowledge has been gained on the non-coding transcriptome and RNAs that are post-transcriptionally derived from them. tRNA-derived small RNAs, which were initially described in bacteria as the 3-dimensional structure of tRNA was being solved, hinted at non-canonical functions by virtue of interaction with other cellular structures^49, 50^. A range of subsequent studies demonstrated that these fragments were generated through a regimented process and implicated several ribonucleases important in their biogenesis^16, 20, 44^. tRNA fragments are classified based on their position relative to the parental molecule and include 5’-tRNA halves or fragments, 3’-tRNA halves or fragments and internal fragments; together, we refer to them as tRNA-derived small RNAs (tDRs). Importantly, several studies have suggested key functional roles for tDRs in the cellular response to stress, affecting fundamental processes such as mRNA stability and silencing, ribosomal function, and stress granule formation^16, 20^. Finally, tDRs appear to constitute a significant proportion of RNAs found in the extracellular compartment across different biofluids, and their export from cells has been shown to affect cellular phenotype or mediate intercellular signaling^51^. However, the presence of modifications on tRNA bases (that may prevent consistent reverse transcription), and the rapidly evolving algorithms for mapping and naming tDRs has been an obstacle in generating a comprehensive atlas of cellular and extracellular tDRs in response to different cellular stressors.

We have previously described methodology that leverages the activity of enzymes that demethylate commonly modified bases in tRNA to better identify previously unreported tDRs^19^. Here we systematically use ARM-seq to comprehensively profile the cellular and extracellular tDR landscape in 4 different cell-types under different cellular perturbations. Our study provides for the first time, a comprehensive atlas of tDR signatures for cellular stress, and surprisingly demonstrates the dynamic changes in extracellular tDRs in response to stress. Finally, in a small pilot study, we demonstrate the possible application of these signatures to human studies; in patients undergoing cardiopulmonary bypass, a model of oxidative stress and nutritional deprivation, we note the dynamic changes in key tDRs previously identified in our cellular studies.

### ARM-seq to identify cellular and extracellular tDRs

As expected from our previous studies, ARM-seq led to a dramatic improvement in the detection of tDRs, with a significant increase in both the proportion of reads mapping to tRNA genes, as well as longer reads that spanned known methylation sites. Most notable was the robust detection of key tDRs in response to cellular stress that were not detected by conventional RNAseq (Fig. S2C). This study therefore represents a far more accurate and comprehensive atlas of the tDR signatures of cellular stress.

This study represents the first use of ARM-seq to detect the profile of extracellular tDRs in response to cellular stress. We chose to examine these profiles across diverse cell lines across two different species (rodent and human) to define whether there are ‘common’ signatures to different cellular stressors. At the same time, we found interesting differences in the proportion of tDRs (both cellular and extracellular) in the cells. For example, CMs appear to have higher tDR expression at baseline and both CMs and CFs show a robust up-regulation of extracellular tDRs upon reoxygenation. Whether different tissues have significant variance in the overall tDR expression at baseline (in the absence of stress) is not clear; furthermore, any functional implication of this finding warrants future investigation. A general theme that emerged when examining the tDRs across the different cells was that the extracellular profile of tDRs in terms of abundance and type diverged significantly from the cellular tDRs: for example, higher proportions of tDRs derived from tRNA-Glu and tRNA-Pro were noted in the extracellular compartment. Our findings complement other published studies^29, 30^, notably the presence of 5’ halves from tRNA-Glu-TTC, tRNA-Glu-CTC, tRNA-Gly-GCC and tRNA-Gly-CCC. However, the use of ARM-seq also allowed for the detection of species other than 5’-halves that have been previously described in the aforementioned studies, such as the 3’-halves derived from tRNA-Asp-GTC, tRNA-Glu-CTC, and tRNA-Asn-GTT, 3’ fragments derived from tRNA-Ser-GCT and tRNA-Pro-AGG, and those without conventional names (e.g. tDR-1:56-Pro-AGG-2-M3-G33A). We recognize that we did not make a concerted effort to separate different ex-RNA carriers (e.g. riboproteins, EVs of different size or density) in these studies; as previously noted, the tDRs we describe may be associated both with EVs or with non-EV carriers as has been previously shown^52^. While several studies have reported that a significant majority of extracellular tDRs are carried in association with riboproteins, other studies have demonstrated the presence of tDRs within EVs^30^, and importantly, a functional role of EV-contained tDRs^53^. Finally, there has been recognition that tDRs from bovine serum used for cell culture may confound analysis of extracellular tDRs; while we have used bovine serum depleted of EVs by ultracentrifugation, we cannot exclude the presence of these confounders. However, as the same medium was also used for the stress conditions, the changes we observed in tDRs cannot be attributed to such confounders.

Like the previously mentioned studies, our work also confirmed the preponderance of 5’-tRNA halves ending in the anti-codon loop, but also demonstrated the presence of 3’-halves and intracellular tDRs that have not been previously recognized in the extracellular compartment. Interestingly, the extracellular tDRs derived from the 3’ halves had distinct ends (at position 74) compared to their cellular counterparts. Whether these findings reflect distinct biogenesis machinery for extracellular tDRs or selective export of these tDRs cannot be inferred from our study. Notably, Nechoostan et al have demonstrated the export of full-length tRNAs into the extracellular non-EV fraction and subsequent processing by RNase 1^32^. The ARM-seq methodology was not designed to detect full length tRNAs, and it is therefore not surprising that we did not detect the parental tRNAs for the abundant tDRs that were detected. Other technologies such as DM-tRNA-seq or the use of may have the ability to tackle this question. Finally, it remains possible that the abundance of certain tDRs may reflect not selective biogenesis, but increased stability of these fragments in the extracellular compartment compared to others^54^. Together our findings suggest that extracellular tDRs have distinct signatures and sequences that do not simply reflect the stoichiometry of cellular tDRs and support the findings of others that the biogenesis of extracellular tDRs may be distinct from cellular tDRs^24, 25, 32^.

### Extracellular tDRs as unique signatures of cellular stress

Our results demonstrated that extracellular tDRs had distinct non-overlapping signatures in response to the different stressors and had far better discrimination between the different stressors than microRNAs. Interestingly while cellular tDRs changed with stress, as has been previously demonstrated, the pattern of expression changes had considerably more overlap between the different stressors than the extracellular tDRs. This pattern was seen in all four cell types examined, suggesting that extracellular tDRs may provide better discrimination in measuring cellular stress than extracellular miRNAs. The functional implications of these findings are not clear. Whether extrusion of tDRs in response to particular stressors leads to changes in cellular phenotypes as has been shown for T-cell activation^55^, or whether cellular stress leads to an upregulation of extracellular tDR biogenesis would of great interest to determine in future studies.

We systematically tracked cellular and extracellular tDRs that were altered in response to each of the stressors (nutrient deprivation, hypoxia or oxidative stress) in each cell line to determine if there were signatures that were common to all cell lines, and if there were tDRs that were unique to individual cell types. When examining nutrient deprivation in the human cell lines, we found far more extracellular tDRs (188 total) that were regulated in the same direction in both HEK and BeWo cells. The top upregulated tDRs were derived from the 5’ halves of tRNA-Gly (which has been previously shown to be a major contributor to EV-contained tDRs^30^), while the top down-regulated tDRs were derived from tRNA-Glu. Similar to the human cell lines, a large number of extracellular tDRs were altered in the rodent CM and CF cells, with the majority being downregulated. Importantly, we found a set of tDRs (Fig. 4H) that were commonly upregulated (tDR-1:36-Gly-CCC-1, tDR-1:35-Gly-CCC-1, tDR-1:36-Asp-GTC-2, tDR-1:36-Glu-CTC-1-M2-C35U, and tDR-3:36-Gly-GCC-1) or down-regulated (approximate 85 tDRs). If validated in multiple other cell-types, these may comprise a ‘universal marker’ for nutrient deprivation or metabolic stress.

Similar findings were noted with hypoxic stress with more pronounced changes in extracellular tDRs, although it appeared that a majority of tDRs were decreased with hypoxic stress. Interestingly, the cancer BeWo cell line showed less pronounced changes than HEK cells in this regard. Whether cancer cells are more resistant to hypoxia, and this underlies these differences would be of interest to address in the future. As compared to these cell lines, the cardiac primary rodent cells had a far higher number of altered extracellular tDRs with a number of these being common between all the cell types.

The generation of tDRs in response to oxidative stress has been shown to be a conserved response in eukaryotic cells. Interestingly, we found some marked differences between the human cell lines (most cellular and extracellular tDRs were decreased) and the primary cardiac cells (where reoxygenation led to an increase in most extracellular tDRs). Notably, the top upregulated tDRs were all derived from the 5’ end of tRNA-Glu-TTC. These data suggested that the cellular context was important in the biogenesis or export of extracellular tDRs in response to oxidative stress.

### Extracellular tDRs as biomarkers for human diseases

The potential of tDRs as biomarkers to diagnose disease or monitor disease progression has been studied to some extent in cancer patients^56^. However, our findings that extracellular tDRs do not necessarily mirror the stoichiometry of cellular tDRs should caution investigators about extrapolating altered levels of tDRs in tissues (such as cancer cells) and expecting similar changes in plasma. Our finding that extracellular tDRs are readily altered with cellular stress led us to query whether similar changes may happen in human subjects, and whether some of the signatures we observed in our cell models were translatable to a clinical context. We examined the plasma tDRs in a pilot study of patients undergoing cardiopulmonary bypass, a model of oxidative stress and nutrient deprivation in human subjects. By measuring tDRs at the initiation (pre-stress) and 2 hours after CPB (peak stress), we could compare the levels of individual tDRs. Interestingly, we found that similar to our cellular experiments, we detected both 5’tDRs and 3’tDRs in plasma with ARM-seq, and that the fragmentation pattern was similar to what was noted in the extracellular tDRs in the cell culture system. We again found that plasma tDRs were able to discriminate between pre-stress and peak-stress samples. Interestingly, 8 out of 17 differentially expressed tDRs after 2 hours of CPB were common with our cellular models of nutrient deprivation, while 5 of these overlapped with signatures of oxidative stress. One of these, tDR-34:75-Met-CAT-1, could only be detected 2 hours after CPB.

It should be noted that this was a preliminary pilot study with a small sample size; however, we were encouraged by our results that some of the tDR signatures of cellular stress we had identified could be readily translated into a clinical context. Whether the presence of these signatures or degree of change is associated with clinical outcomes would be of paramount importance in future studies.

## Conclusion

Using ARM-seq we have provided a comprehensive profile of cellular and extracellular tDRs in response to various cellular stressors in several different cell types. Our studies demonstrate that the profile of extracellular tDRs and their biogenesis may differ from cellular tDRs, and that the response to stress leads to robust changes in the extracellular tDRs. Notably, extracellular tDRs provide far better discrimination between different types of cellular stressors than miRNAs. In a pilot study, we demonstrate applicability of our findings in a human subject model of nutrient deprivation/oxidative stress. We expect that our data may be of use to investigators in this field who wish to investigate extracellular tDRs as biomarkers for disease processes, or examine their functional role in the context of cellular stress.

## METHODS AND MATERIALS

### Recombinant His-Alkb Expression and Purification

The expression and purification of recombinant His-AlkB were performed as previously described with modifications^57^. Briefly, the His-AlkB-AVA421 plasmid was transformed into *E. coli* BL21(DE3)pLysS competent cells (Promega, Cat# L1191) and cultured in Overnight Express™ Instant TB Medium (Sigma, Cat# 71491) at 37 ℃ to OD^600^ 1.2. The cell pellet was resuspended and lysed by sonication and the cell debris was removed by centrifugation. Then, the clarified supernatant from the crude extract was carefully collected and mixed with TALON Metal Affinity Resin (Takara Bio, Cat# 635502) for 1 hour to allow for the binding of His-tagged AlkB protein to resin. After several washing steps, the resin-bind proteins were eluted and separated by Econo-Column® Chromatography Columns (BioRad, Cat# 7371512). The protein concentration of eight collected fractions was measured and the three protein-enriched fractions were pooled and dialyzed successively in Dialysis Buffer and Storage Buffer. The dialyzed protein supernatant containing recombinant His-AlkB protein was validated by Coomassie Bule staining and western blotting, and used for ARM-seq.

### Human Cell Line Culture and Stress Treatment

HEK293 cells were purchased from ATCC (ATCC, CRL-1573) and cultured in D10 medium, which consisted of DMEM with high glucose and pyruvate (Thermo Fisher, Cat# 11995073) supplemented with 10% fetal bovine serum (FBS) (Thermo Fisher, Cat# 10437028) and 1% Penicillin/Streptomycin (Thermo Fisher, Cat# 15140122). BeWo cells (ATCC, CCL-98), were cultured in F-12K Medium (ATCC, Cat# 30-2004) containing 10% FBS and 1% Penicillin/Streptomycin. For the glucose and serum deprivation (GSD) treatment, HEK293 and BeWo cells were cultured in glucose-free DMEM basal medium for 24 hours. For the hypoxia treatment, HEK293 and BeWo cells were fed with complete medium and maintained in hypoxia chamber with 0.1% oxygen for 24 hours. For the oxidative stress treatment, HEK293 and BeWo cells were treated with 0.8 mM hydrogen peroxide (H_2_O_2_) and 1.7 mM H_2_O_2_ separately in complete medium for 24 hours. FBS used for control, hypoxia and H_2_O_2_ treatments was filtered through 0.2 μm filter and followed by 24 h ultracentrifugation at 120,000 g to deplete extracellular vesicles (EVs) before use.

### Neonatal Rat Ventricular Cardiomyocytes (CMs) and Cardiac Fibroblast (CFs) Isolation and Stress Treatment

All animal procedures conformed to the animal welfare regulations of the Massachusetts General Hospital Subcommittee on Research Animal Care (SRAC). Neonatal rat ventricular CMs and CFs were isolated as previously described^38^. Briefly, the ventricular heart tissues were carefully dissected from 1-day old Sprague Dawley rat neonates and minced using a sterile sharp razor blade, followed by 8 rounds of digestion in the ADS buffer (NaCl 116 mM, HEPES 20 mM, NaH_2_PO_4_ 1 mM, KCl 5.4 mM, MgSO_4_ 0.8 mM, and Glucose 5.5 mM) containing 0.05 mg/ml collagenase type II (Worthington, Cat# LS004177) and 1 mg/ml pancreatin (Sigma, Cat# P3292). Then the single cell suspensions were pooled and plated into 10 cm tissue culture dishes with CM medium [DMEM medium supplemented with 10% horse serum (Thermo fisher, Cat# 26050-088), 5% FBS and 1% Penicillin/Streptomycin] for 1 hour. The attached cells were considered to be CFs and maintained in D10 medium. The floating cells were then purified via Percoll (GE Healthcare, Cat# 17-0891-01) gradient; the cells in the middle layer, which were considered as CMs, were collected and maintained in CM medium.

For the GSD treatment, CF and CM cells were exposed to glucose-free DMEM basal medium for 5 hours. For the hypoxia treatment, CF and CM cells were fed with glucose free DMEM basal medium and maintained in hypoxia chamber with 0.2% oxygen for 5 hours. For the reoxygenation (ReO_2_) treatment, CF and CM cells were exposed to hypoxia treatment for 5 hours and then fed with complete medium and cultured in normoxia for an additional 24 hours. FBS and horse serum used for ReO_2_ treatment were also EV-depleted by filtration and ultracentrifugation.

### CPB Patient Plasma Sample Collection

Plasma samples were prospectively collected from six patients undergoing elective aortic valve replacement surgery with cardiopulmonary bypass by a single surgeon at a single institution in 2020 (https://clinicaltrials.gov/ct2/show/NCT00985049). Written informed consent approved by the Partners Healthcare Institutional Review Board (Boston, MA) was provided. Baseline plasma samples were obtained from venous blood immediately after commencement of cardiopulmonary bypass, which included intermittent cold blood cardioplegia for myocardial protection. Post-ischemic samples were obtained immediately before removal of the aortic cross-clamp at the end of the de-airing procedure. Plasma samples were stored at −80℃ before RNA isolation. Median ischemic time was 73 minutes (Inter-quartile range [IQR] 68-121). Of the six patients included in this analysis, 3 (50%) were female and the median age was 73 years (IQR 66-75).

### Cellular Small RNA and Extracellular RNA Isolation

Cells were immediately lysed by adding TRIzol™ (Thermo Fisher, Cat# A33251) after treatment and total RNAs were then isolated by following the manual. 40 μg total RNAs were subjected to small RNA isolation by using mirVana™ miRNA Isolation Kit (Thermo Fisher, Cat# AM1560). The cell culture medium was collected immediately after treatment and spun twice at 2,000 g for 10 min to remove cell debris, followed by filtering through 0.8 μm filter. Next, 10∼12 ml cell culture medium was subjected to extracellular RNA isolation using the exoRNeasy Maxi kit (Qiagen, Cat# 77164) with minor modifications. 20 μg/ml Glycogen was added into the upper aqueous phase before adding 2 volumes of 100% ethanol and the aqueous phase/ethanol mixture was incubated at -20℃ overnight to facilitate with small RNA precipitation. The extracellular RNAs from 1 mL CPB patient plasma samples were isolated using the exoRNeasy Midi kit (Qiagen, Cat# 77044).

### Quantitative Polymerase Chain Reaction (qPCR)

qPCR was performed as previously described^58^. Briefly, 1 μg of cellular total RNA was reverse transcribed to cDNA using the High-Capacity cDNA Reverse Transcription Kit (Thermo Fisher, Cat# 4368813). qPCR was then performed using a QuantStudio 6 Flex Real-Time PCR Systems (Thermo Fisher) with SsoAdvanced™ Universal SYBR® Green Supermix (BioRad, Cat# 1725275). The sequences of qPCR primers used are listed in Table S7.

### Western Blot

Western blotting was performed as previously described^58^. Briefly, cell pellets were resuspended in RIPA buffer (Thermo Fisher, Cat# 89900) containing Protease and Phosphatase Inhibitor Cocktail (Thermo Fisher, Cat# 78441), and PMSF, and lysed via sonication. The protein concentration of clarified cell lysate was then measured using the BCA protein assay kit (Thermo Fisher, Cat# 23227). 5 ∼ 20 μg of total protein per lane was separated by SDS-PAGE gel and transferred to PVDF membrane. 5% BSA in TBST was used for blocking and primary antibody incubation. HRP conjugated secondary antibodies were then applied and the blots were developed using SuperSignal® West Femto Maximum Sensitivity Chemiluminescent Substrate (Thermo Fisher, Cat# 34094). The antibodies used were the following: HIF1a (Cell Signaling Technology, Cat# 14179S), DDIT3 (Cell Signaling Technology, Cat# 2895), DDIT4 (Proteintech, Cat# 10638-1-AP), r-H2AX (Cell Signaling Technology, Cat# 80312), LC3B (Cell Signaling Technology, Cat# 3868S), β-Actin (Sigma, Cat# A5316), Goat Anti-Mouse Immunoglobulins/HRP (Dako, Cat# P0447), and Goat Anti-Rabbit Immunoglobulins/HRP (Dako, Cat# P0448).

### RNA Sample Pre-treatment and Small RNA Library Preparation for ARM-Seq

Cellular small RNAs and extracellular RNAs were successively treated with DNase I (Zymo Research, Cat# R1014), T4 Polynucleotide Kinase (New England Biolab, Cat# M0201L), and His-AlkB. After each treatment, the RNA was cleaned to remove enzyme and buffer components using the RNA Clean & Concentrator-5 kit (Zymo Research, Cat# R1014) before the next treatment, with the following minor modifications. 20 μg/ml Glycogen was added into sample/RNA Binding Buffer mixture before adding an equal volume of 100% ethanol and then the mixture incubated at -20℃ overnight to facilitate small RNA precipitation. CPB patient plasma RNAs were pretreated with Heparinase I (New England Biolab, Cat# P0735L) at 30 ℃ for 3 hours before DNase I treatment. Then, 100-500 ng His-AlkB treated RNAs were used as input into the NEBNext Multiplex Small RNA Library construction kit (New England Biolab, Cat# E7300S) and instructions were followed with the exception of 60 ℃ temperature used for reverse transcription step. Size selection was performed by isolating the bands from 140 bp – 240 bp to remove the contamination of primer adaptors (around 127 bp) and longer RNAs. The quality of libraries was confirmed by High Sensitivity DNA ScreenTape Analysis (TapeStation, Agilent) and qPCR. Multiplexed libraries were sequenced on an Illumina® HiSeq 2000 system with paired-end 75 bp sequencing. About seven million read pairs were generated from each library.

### ARM-seq Data Processing

#### Mapping and tDR naming

The accession number for the raw sequence data of in vitro stress response platforms reported in this paper is GEO: GSE173806. Paired-end sequencing reads were trimmed of their adapter sequences and merged using SeqPrep with the following parameters: -L 15 -A AGATCGGAAGAGCACACGTC -B GATCGTCGGACTGTAGAACTC. Reads were aligned to the pre-built tRAX [http://trna.ucsc.edu/tRAX/] hg38 or rn6 reference database using Bowtie2 in very-sensitive mode with the following parameters to allow for a maximum of 100 alignments per read: --very-sensitive --ignore-quals --np 5 -k 100. Utilizing the tRAX pipeline [https://github.com/UCSC-LoweLab/tRAX], ENSEMBL (release 96) small ncRNA annotations were used to count the number of small RNA transcripts using only primary alignments to prevent double counting. For tDRs, tDRnamer [http://trna.ucsc.edu/tDRnamer/] was used to generate unique identifiers for each of the transcripts based on the isodecoder(s) they mapped to and positions of any misincorporations. The naming conventions for tDRnamer can be found at http://trna.ucsc.edu/tDRnamer/docs/naming/, and are based on the full-length tRNA names derived from the Genomic tRNA Database^59^. Raw counts were normalized using the DESeq2 package and utilized for downstream analyses.

#### Read length distribution

To determine the read length distribution for different small RNA types reads were extracted that either aligned to a tRNA feature or another small RNA feature deemed “other”. The proportion of read lengths was then calculated by taking the number of reads corresponding to each length and dividing that by the total number of reads for that category (tRNA or other).

#### tDR abundance and end distribution

To determine the abundance distribution for tDRs across a consensus tRNA transcript, tDR normalized read counts were extracted. For each of these transcripts, tRNA mapping information was used to determine which positions in the mature tRNA transcript these tDRs were derived from. Using this information, all tRNAs were collapsed into a “consensus” tRNA and the proportion of total abundance at each position was calculated by dividing the total abundance at each position in the consensus tRNA by the total number of tDR reads.

To determine the read end distribution for these tDRs across a consensus tRNA transcript, abundance values were attributed to only the final position the tDR mapped within the mature tRNA transcript. The end frequency was then calculated by dividing the total abundance at each position in the consensus tRNA by the total number of tDR reads.

#### Dimensionality reduction and correlations

Using DESeq2 normalized values, dimensionality reduction was performed to assess the reproducibility of biological replicates. These analyses were performed for all tDRs and miRNAs, as annotated by ENSEMBL (release 96) by first extracting read counts for those transcripts prior to dimensionality reduction. Principle component analysis (PCA) was performed using the Python package sklearn v0.23.2 and the first two components were used to generate scatter plots. UMAP projections for each of the small RNA types were generated using the Python package umap-learn v0.5.0. Spearman correlation coefficients were calculated using the pandas v1.2.1 “corr” function with parameters: method=’spearman’.

### Tracking plots

Tracking plots were generated to track the abundance of specific tDRs and miRNAs across different cellular conditions and/or environments using DESeq2 normalized values. Initially, tDRs were filtered to remove potential sequencing artifacts and degradation products by setting a cutoff threshold across all replicates. For miRNAs, all transcripts with >5 reads were retained. Biological replicates were merged and log2 values were calculated for all transcript types.

### Statistical Analysis

Statistical analyses of qPCR and patient sequencing data were performed using GraphPad Prism software (version 9). qPCR data is expressed as mean ± SEM and the statistical significance was assessed by two-tailed unpaired Student’s t test; for CPB patient ARM-seq data, two-tailed paired Student’s t test was used; for all other ARM-seq data, differential expression analysis was performed using DESeq2 to generate Benjamini–Hochberg-corrected P values (padj) to assess the statistical significance. The criterion for statistical significance was P < 0.05 (* P < 0.05, ** P < 0.01, *** P < 0.001).

## ACKNOWLEDGEMENTS

G.L. is supported by an American Heart Association (AHA) postdoctoral fellowship (19POST34381027). H.L is supported by a career development grant from AHA (20CDA35310184). J.D.M is supported by grants from NIH (R01HL118266, 1R01HL150401). S.D. is supported by grants from NIH (1R01HL150401, R35HL150807) as well as from NCATS (1UG3 TR002878).

