## Supplemental Methods and Figures 1-14 for "Distinct stress-dependent signatures of cellular and extracellular tRNA-derived small RNAs (tDRs)"

#### Standardized tDR naming system

tDRnamer (http://trna.ucsc.edu/tDRnamer/) was used for the standardized tDR naming in this study, which consists of five parts as shown in the example below:

tDR-4:33-Val-AAC-1-M7-A10U

① ② ③ ④ ⑤

The instruction manual for tDRnamer can be found at http://trna.ucsc.edu/tDRnamer/docs/naming/. Briefly, ①. Prefix - "tDR" stands for "tRNA Derived RNA". ②. Position – This includes the start and end positions of the tDR relative to the source tRNA. If the position is located at the leader or trailer sequence of the precursor tRNA gene, the numbering will be preceded with a letter "L" or "T" respectively. ③. Source tRNA - This is the name of the tRNA from which the tDR is derived. tRNA names from Genomic tRNA Database are used. ④. Matching tRNA transcripts – If a tDR is mapped to multiple tRNA genes, an optional component with prefix "M" and the number of matching tRNAs will be added to the tDR name. ⑤. Variations – For a substitution, the annotation will include the base in tDR, the position of the substitution relative to the tDR, and the substituted base in the source tRNA. For an insertion, the annotation will include a prefix "I", followed by the position of the insertion relative to the tDR and the inserted base. For a deletion, the annotation will include a prefix "D", followed by the position before the deletion relative to the tDR and the deleted base from the source tRNA.

### SUPPLEMENTAL FIGURES

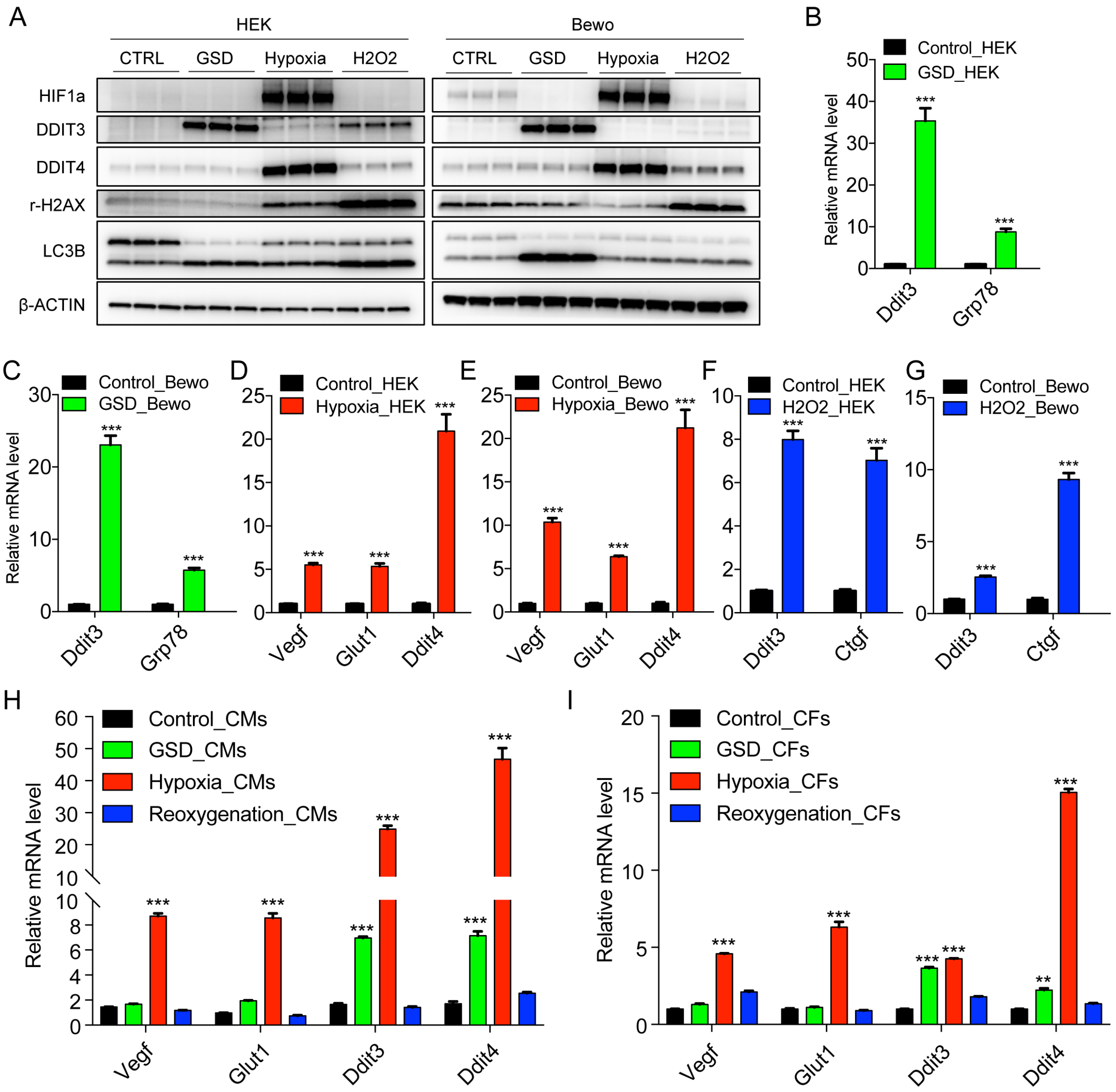


**Figure S1. The validations of *in vitro* stress response platforms**

**A**. Western blot validation of stress response in HEK and BeWo cells.

**B** and **C**. Nutritional deprivation induces Ddit3 and Grp78 expression in HEK cells (B) and BeWo cells (C).

**D** and **E**. Hypoxia activates the expression of Ddit4 and angiogenesis-related genes, including Vegf and Glut1, in HEK (D) and BeWo (E) cells.

**F** and **G**. $H_2O_2$ treatment increases the expression levels of Ddit4 and Ctgf in HEK (F) and BeWo cells (G).

**H** and **I**. QPCR validation of neonatal rat primary CM (H) and CF (I) stress response platforms. Data are shown as means ± SEM of at least three independent experiments. The unpaired two-tailed Student's t test was used in (B) - (I). $^{**}p < 0.01$, $^{***}p < 0.001$ versus the control group.

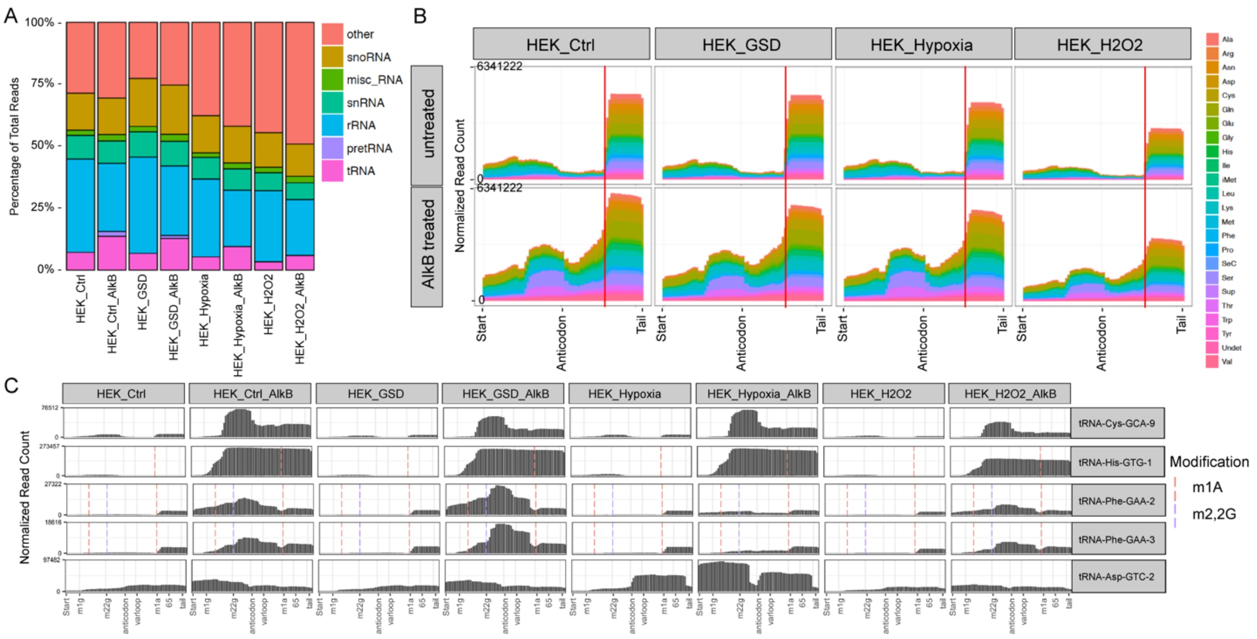


**Figure S2. AlkB treatment increases the abundance and diversity of tRNA reads**

**A**. AlkB treatment dramatically increases the proportion of tRNA reads.

**B**. AlkB treatment significantly elongates the read length of tRNA genes; red line indicates the m1A58 position.

**C**. tRNA coverage plots show four representative tRNAs that could only be detected by ARM-seq but not regular small RNA-seq.

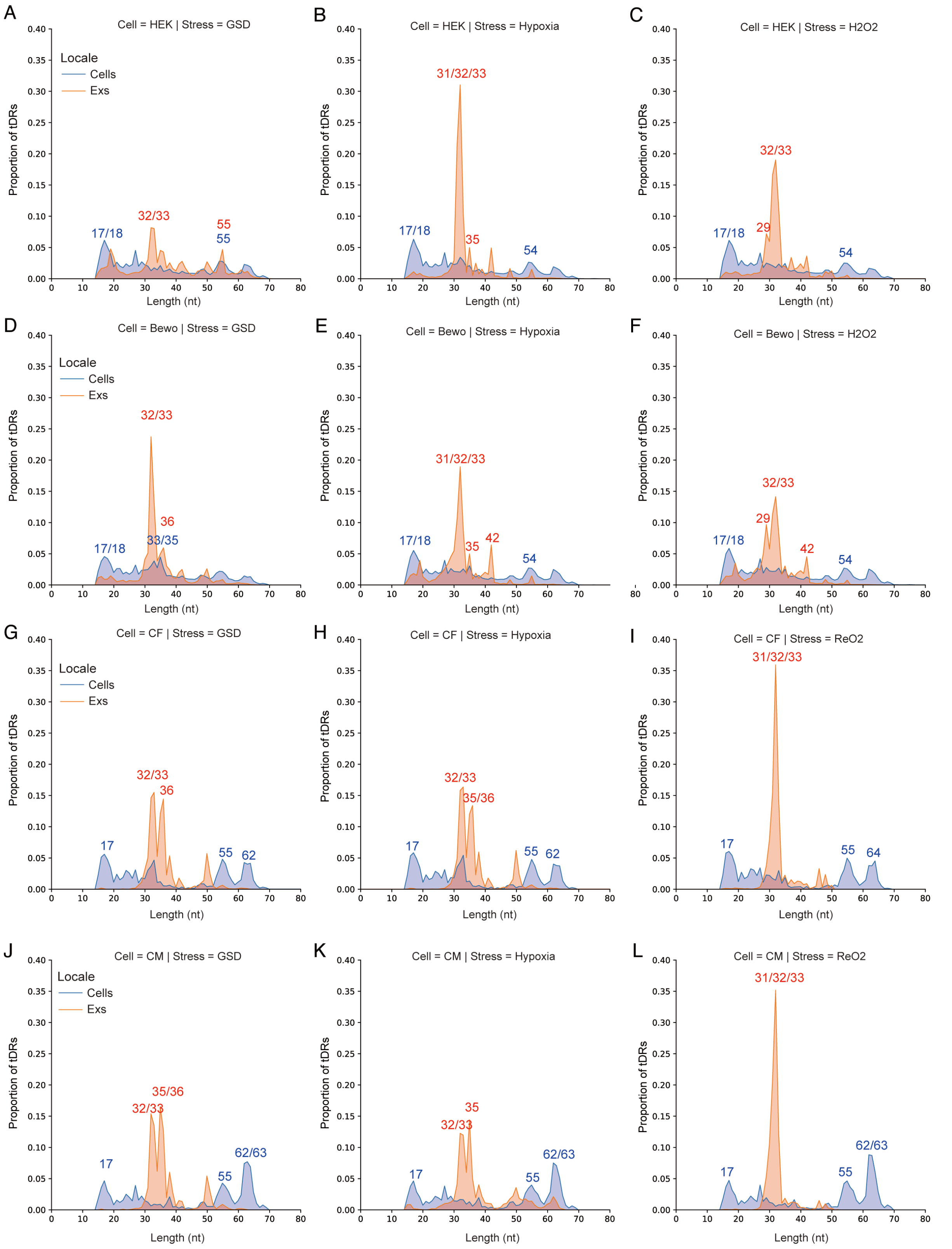


**Figure S3. Extracellular tDRs are predominantly 31-35 nts length in both human and rat samples upon different stress treatments.**

**A-C**. The length distribution plots of intracellular tDRs and extracellular tDRs during GSD (A), hypoxia (B) and $H_2O_2$ (C) treatment in HEK cells.

**D-F**. The length distribution plots of intracellular tDRs and extracellular tDRs during GSD (D), hypoxia (E) and $H_2O_2$ (F) treatment in BeWo cells.

**G-I**. The length distribution plots of intracellular tDRs and extracellular tDRs during GSD (G), hypoxia (H) and $ReO_2$ (I) treatment in CF cells.

**J-L**. The length distribution plots of intracellular tDRs and extracellular tDRs during GSD (J), hypoxia (K) and $ReO_2$ (L) treatment in CM cells.

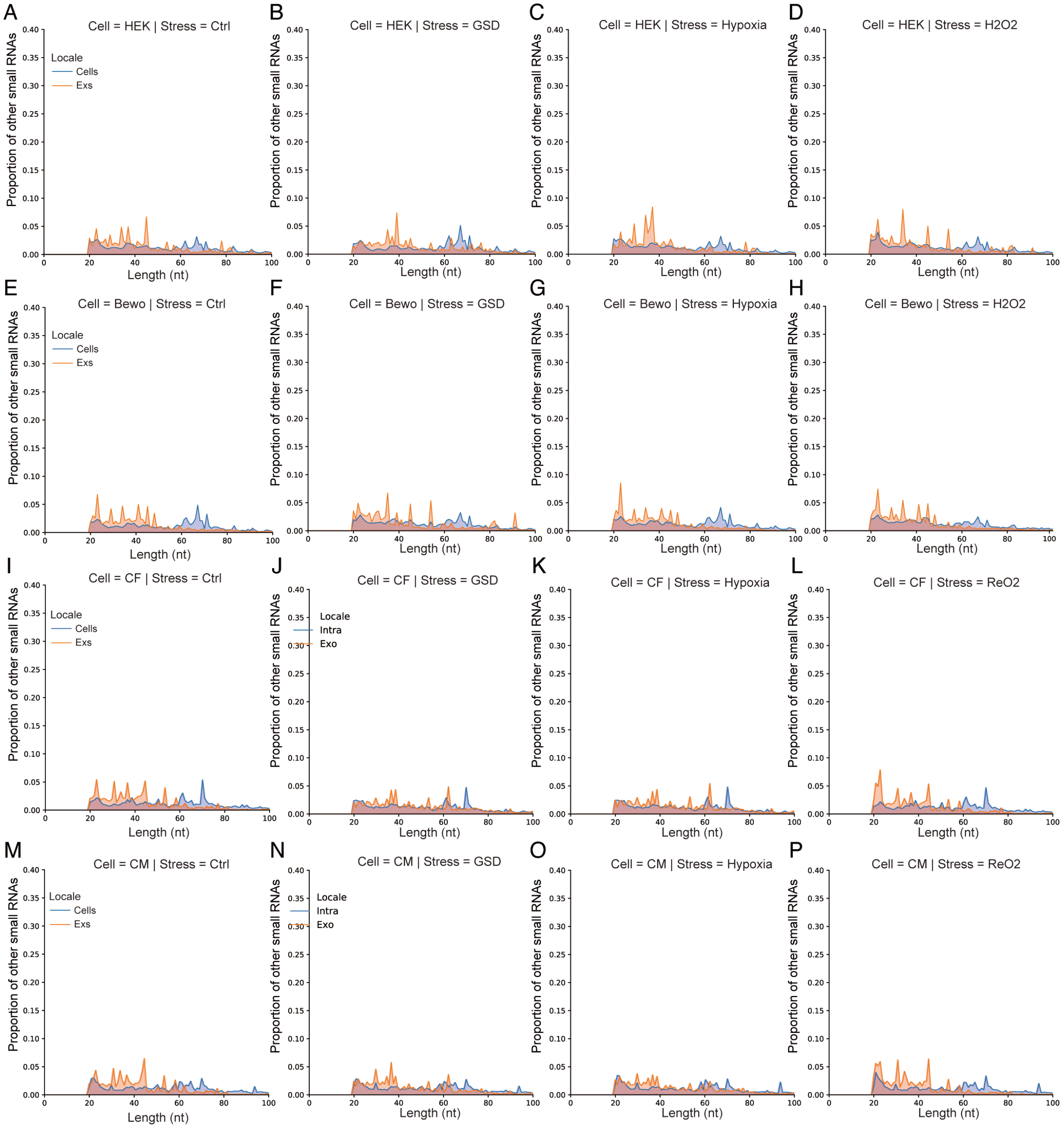


**Figure S4. Other extracellular small RNA species don't have obvious length preference.**

**A-D**. The length distribution plots of intracellular and extracellular other small RNA species at baseline (A) and during GSD (B), hypoxia (C) and $H_2O_2$ (D) treatment in HEK cells.

**E-H**. The length distribution plots of intracellular and extracellular other small RNA species at baseline (E) and during GSD (F), hypoxia (G) and $H_2O_2$ (H) treatment in BeWo cells. **I-L**. The length distribution plots of intracellular and extracellular other small RNA species at baseline (I) and during GSD (J), hypoxia (K) and $ReO_2$ (L) treatment in CF cells. **M-P**. The length distribution plots of intracellular and extracellular other small RNA species at baseline (M) and during GSD (N), hypoxia (O) and $ReO_2$ (P) treatment in CM cells.

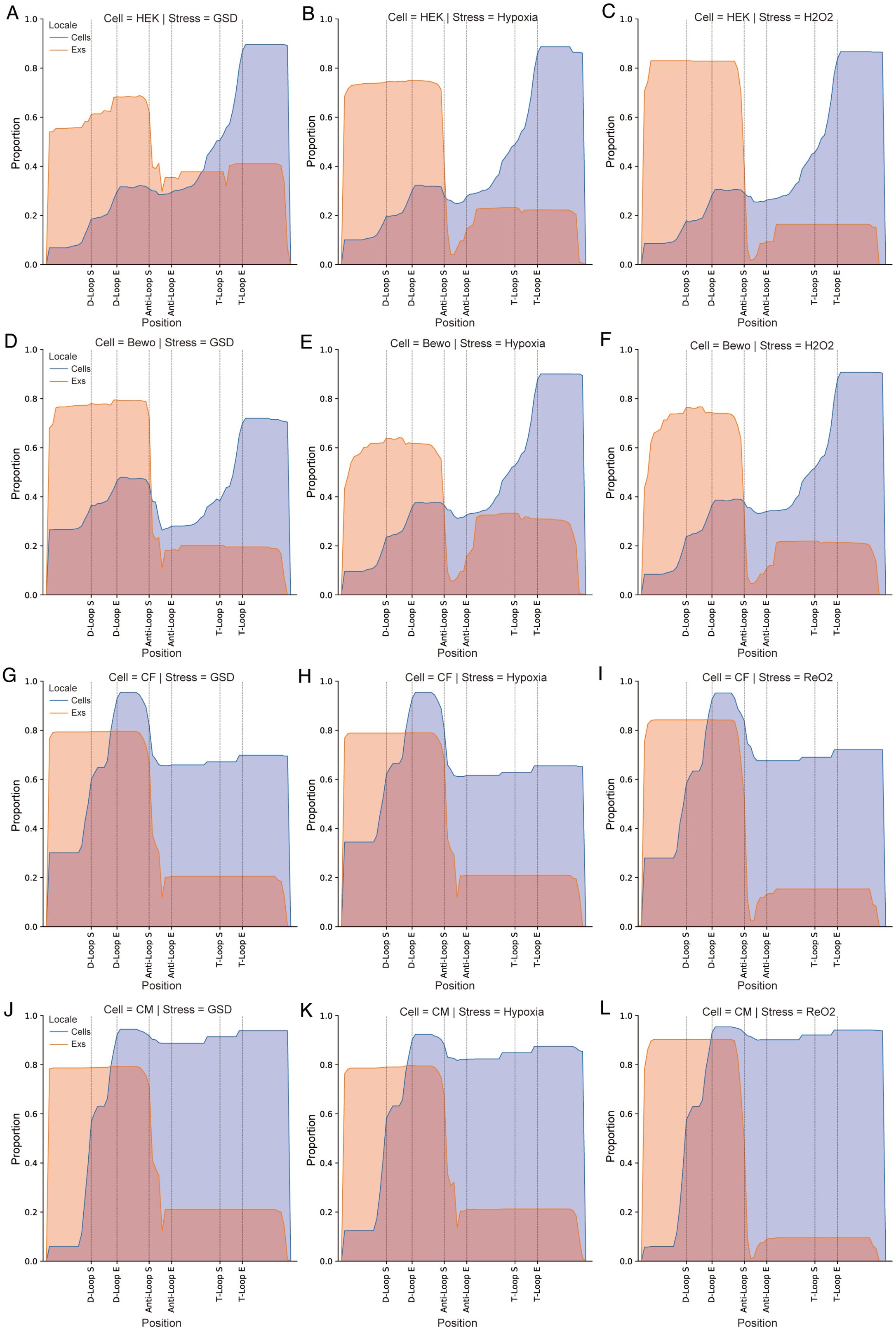


**Figure S5. Extracellular tDRs are mainly derived from 5’ end of tRNA while**

**intracellular tDRs are predominantly derived from 3' end of tRNAs.**

**A-C**. The coverage plots of intracellular tDRs and extracellular tDRs during GSD (A), hypoxia (B) and $H_2O_2$ (C) treatment in HEK cells.

**D-F**. The coverage plots of intracellular tDRs and extracellular tDRs during GSD (D), hypoxia (E) and $H_2O_2$ (F) treatment in BeWo cells.

**G-I**. The coverage plots of intracellular tDRs and extracellular tDRs during GSD (G), hypoxia (H) and $ReO_2$ (I) treatment in CF cells.

**J-L**. The coverage plots of intracellular tDRs and extracellular tDRs during GSD (J), hypoxia (K) and $ReO_2$ (L) treatment in CM cells.

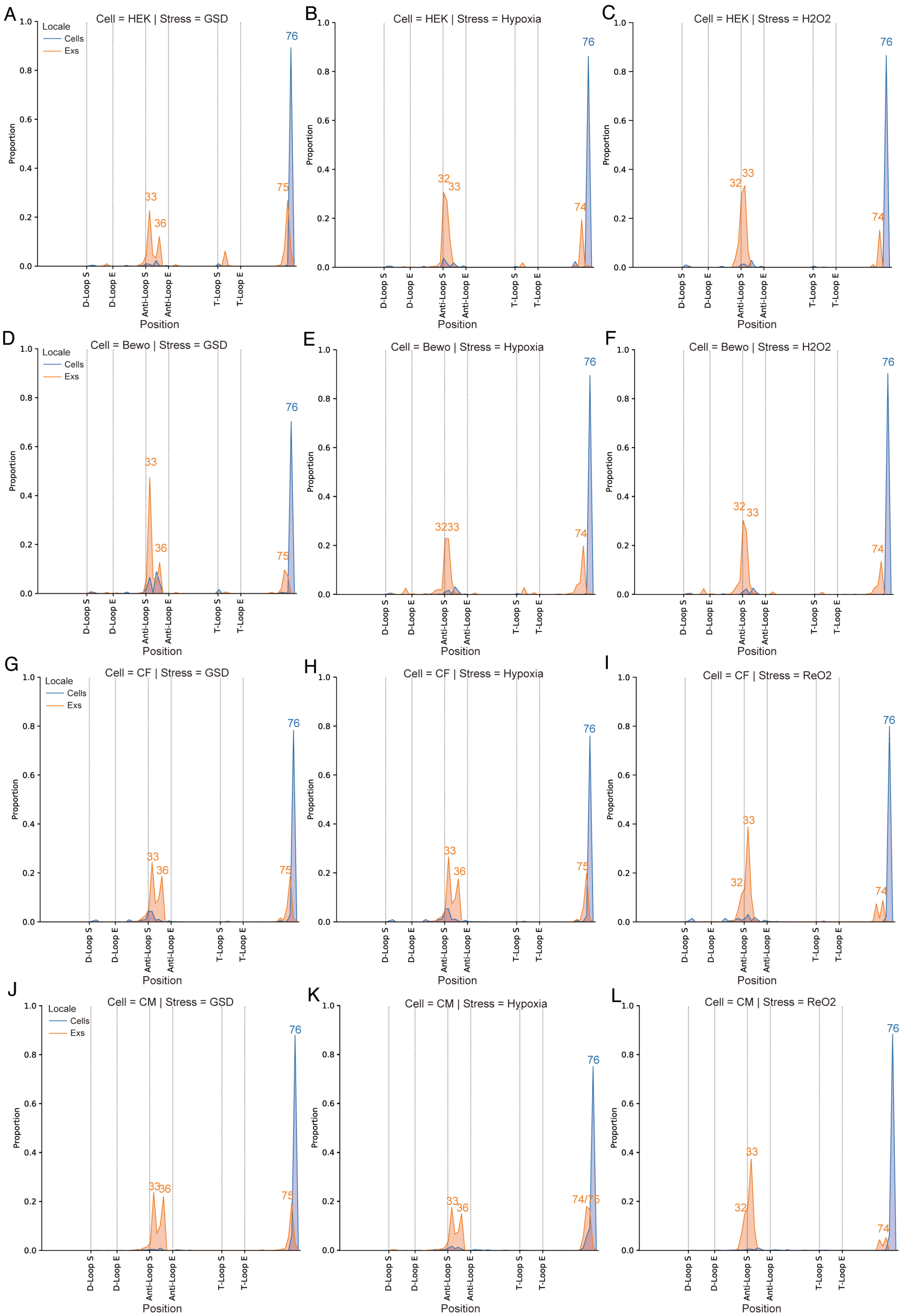


**Figure S6. Extracellular tDRs mainly terminate at either the anticodon loop or position 74 of tRNA genes while intracellular tDRs predominantly terminate at**

**position 76 of tRNA genes.**

**A-C**. The end positions of intracellular tDRs and extracellular tDRs during GSD (A), hypoxia (B) and $H_2O_2$ (C) treatment in HEK cells.

**D-F**. The end positions of intracellular tDRs and extracellular tDRs during GSD (D), hypoxia (E) and $H_2O_2$ (F) treatment in BeWo cells.

**G-I**. The end positions of intracellular tDRs and extracellular tDRs during GSD (G), hypoxia (H) and $ReO_2$ (I) treatment in CF cells.

**J-L**. The end positions of intracellular tDRs and extracellular tDRs during GSD(J), hypoxia (K) and $ReO_2$ (L) treatment in CM cells.

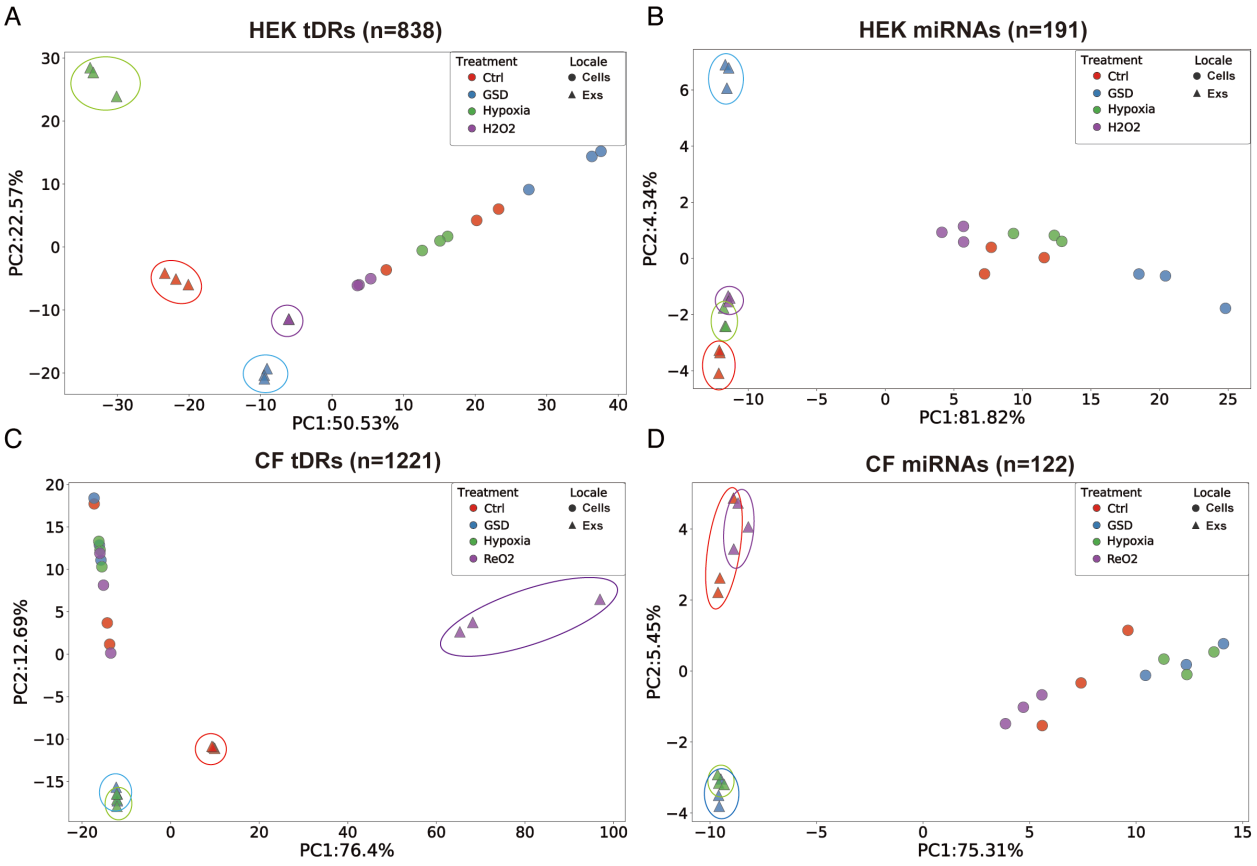


**Figure S7. PCA analysis based on tDR profiles gets better resolution to istinguish different stress responses than those based on miRNA expressions.**

**A** and **B**. PCA analysis of expression values for tDR class (A) provides better resolution to distinguish the Exs samples (circled) derived from HEK cells after different stress treatments than miRNA class (B).

**C** and **D**. PCA analysis of expression values for tDR class (C) provides better resolution to distinguish the Exs samples (circled) derived from CF cells after different stress treatments than miRNA class (D).

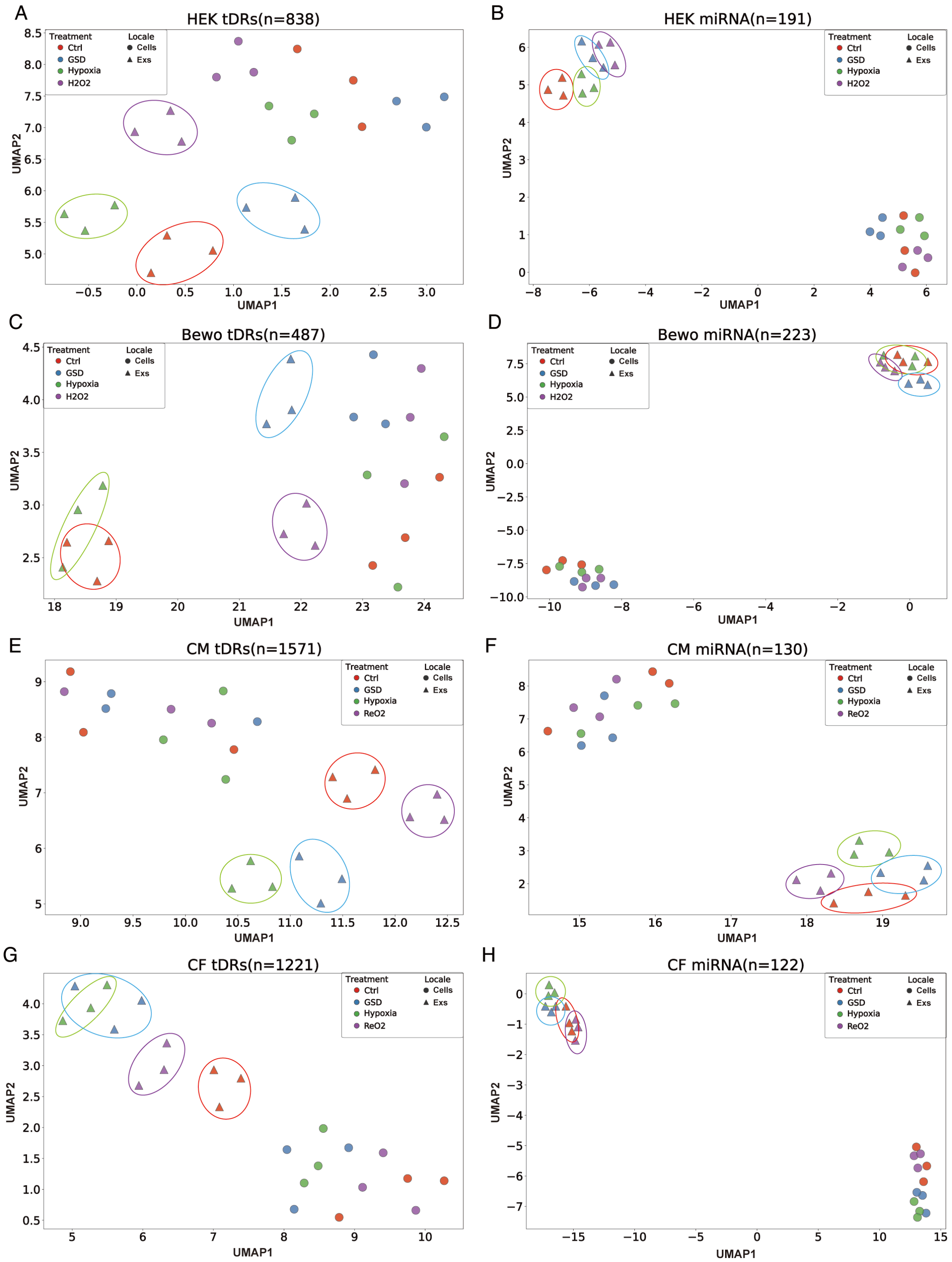


**Figure S8. UMAP projection analysis shows that extracellular tDR signatures provide better resolution to distinguish different stress responses than miRNA**

**signatures**

**A** and **B**. UMAP projection of expression values for tDR class (A) provides better resolution to distinguish different stress-specific Exs samples (circled) from HEK cells than miRNA class (B).

**C** and **D**. UMAP projection of expression values for tDR class (C) provides better resolution to distinguish different stress-specific Exs samples (circled) from BeWo cells than miRNA class (D).

**E** and **F**. UMAP projection of expression values for tDR class (E) provides better resolution to distinguish different stress-specific Exs samples (circled) from CF cells than miRNA class (F).

**G** and **H**. UMAP projection of expression values for tDR class (G) provides better resolution to distinguish different stress-specific Exs samples (circled) from CM cells than miRNA class (H).

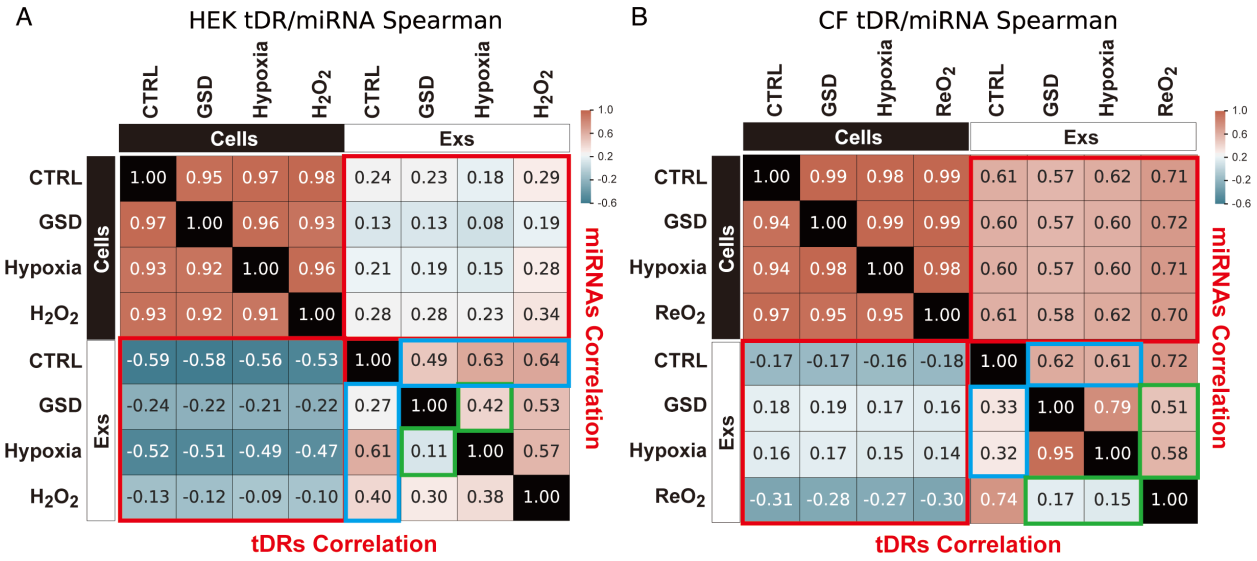


**Figure S9. Extracellular tDR expression landscapes show larger variance among different stress-treated samples than miRNAs**

**A.** Heatmaps of correlation coefficients (Spearman) for tDR class (left bottom) shows larger variance among different samples than miRNA class (right top) in HEK cells.

**B**. Heatmaps of correlation coefficients (Spearman) for tDR class (left bottom) shows larger variance among different samples than miRNA class (right top) in CF cells.

Red boxes show the difference between intracellular samples and extracellular samples; blue boxes indicate the difference of extracellular samples between each stressor and control group; green boxes show the difference between different stressors.

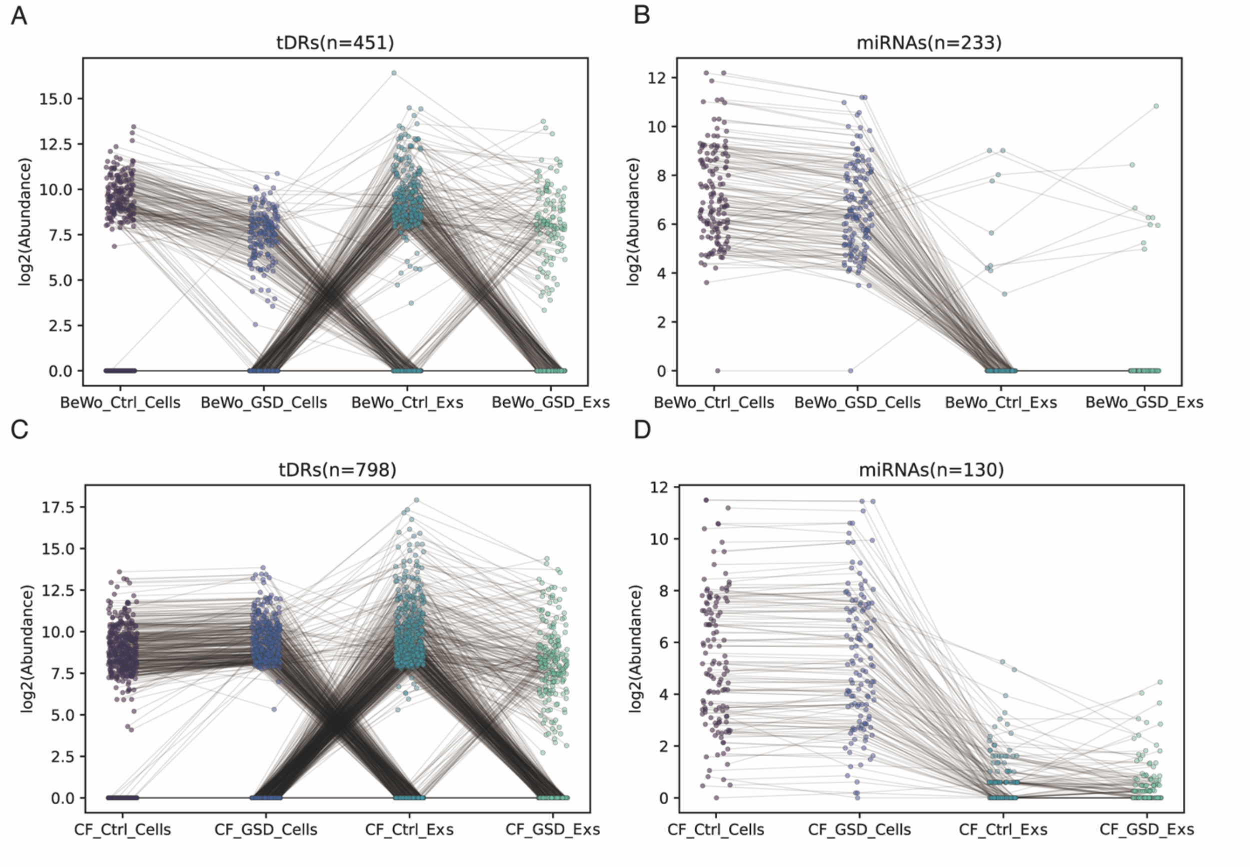


**Figure S10. Nutritional deprivation-shaped dynamic expression of cellular and extracellular tDRs and miRNAs**

**A** and **B**. Expression tracing plots reveal more robust changes in tDR expression (A) when compared with miRNA expression (B) in BeWo cells during GSD treatment;

**C** and **D**. Expression tracing plots reveal more dynamic changes in tDR expression (C) when compared with miRNA expression (D) in CF cells during GSD treatment.

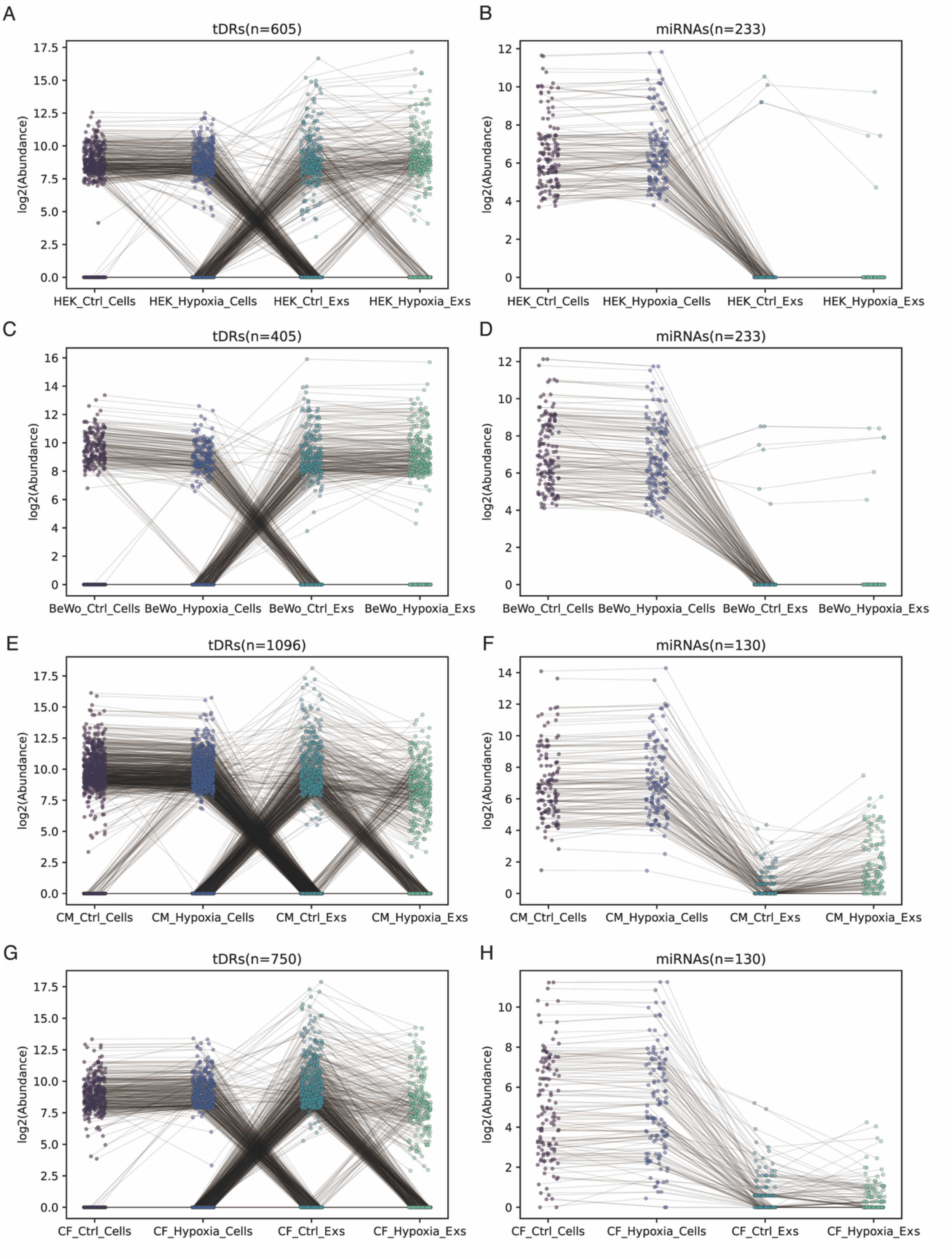


**Figure S11. Hypoxia-shaped dynamic expression of cellular and extracellular tDRs**

**and miRNAs**

**A** and **B**. Expression tracking plots reveal more robust changes in tDR expression (A) when compared with miRNA expression (B) in HEK cells during hypoxia treatment;

**C** and **D**. Expression tracking plots reveal more robust changes in tDR expression (C) when compared with miRNA expression (D) in BeWo cells during hypoxia treatment;

**E** and **F**. Expression tracking plots reveal more robust changes in tDR expression (E) when compared with miRNA expression (F) in CM cells during hypoxia treatment;

**G** and **H**. Expression tracking plots reveal more dynamic changes in tDR expression (G) when compared with miRNA expression (H) in CF cells during hypoxia treatment.

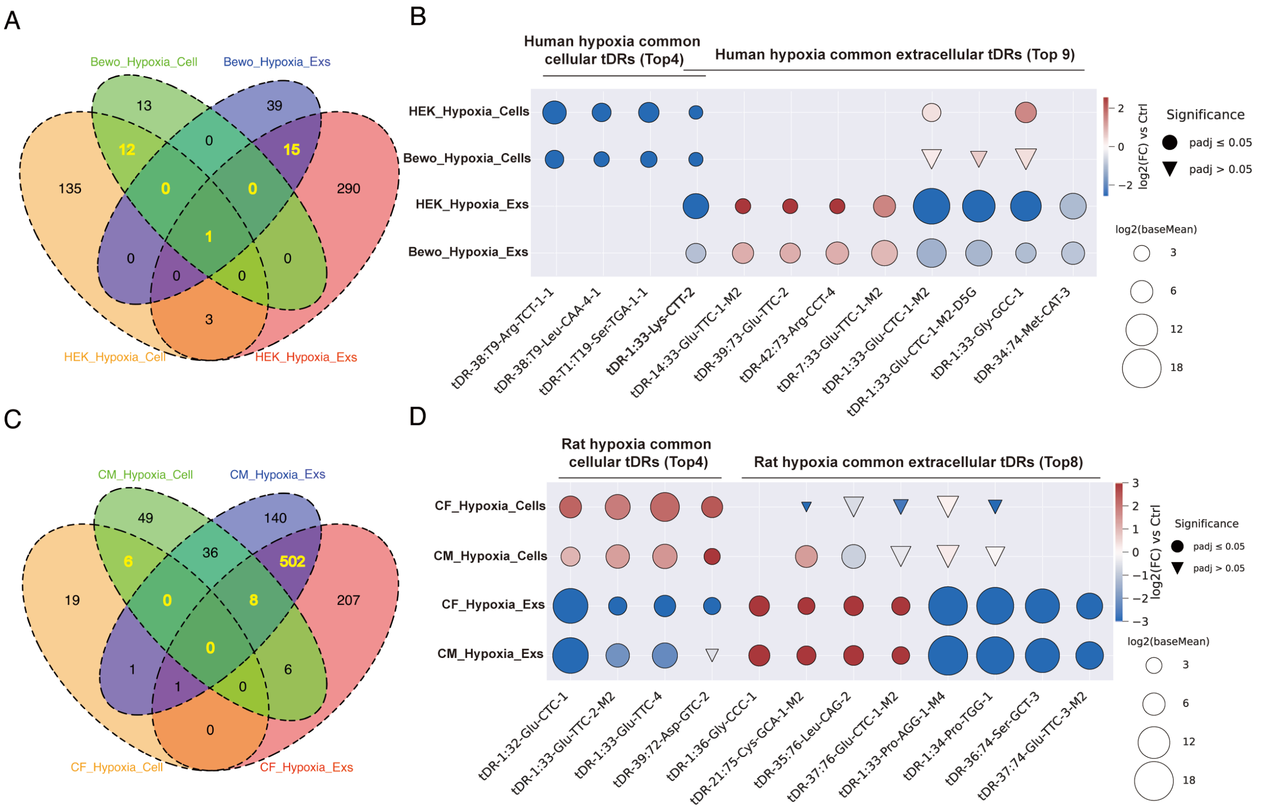

**Figure S12. Hypoxia-shaped cellular and extracellular tDR signatures**

**A**. Venn diagram shows the overlapped and specific tDRs that were regulated by hypoxia in human cells and Exs.

**B**. The most significant cellular and extracellular tDRs that were regulated by hypoxia in both HEK and BeWo cells.

**C**. Venn diagram shows the overlapped and specific tDRs that were regulated by hypoxia in rat cells and Exs.

**D**. The most significant cellular and extracellular tDRs that were regulated by hypoxia in both CF and CM cells.

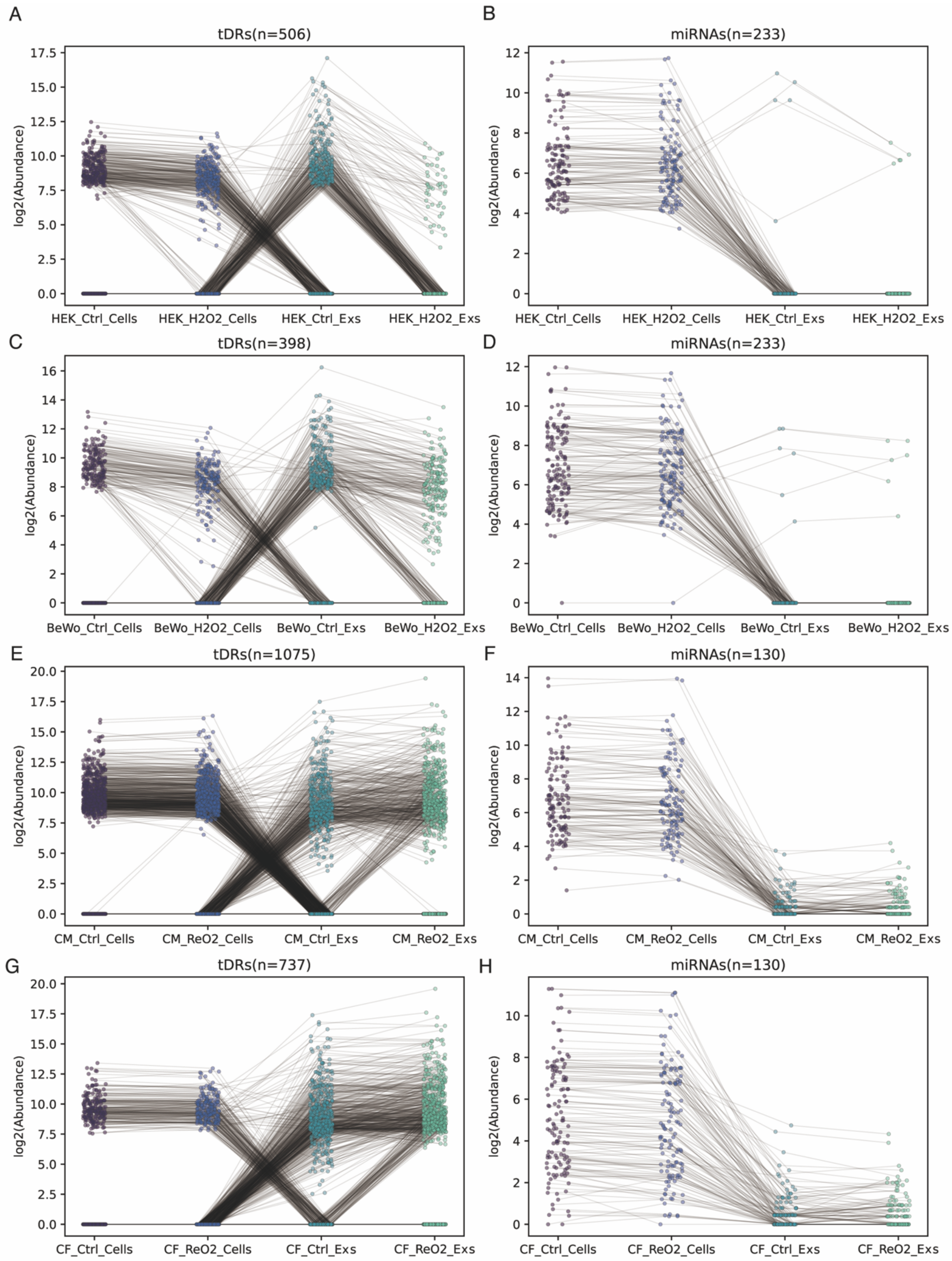


**Figure S13. Oxidative stress-shaped dynamic expression of cellular and extracellular tDRs and miRNAs**

**A** and **B**. Expression tracking plots reveal more robust changes in tDR expression (A) when compared with miRNA expression (B) in HEK cells during $H_2O_2$ treatment;

**C** and **D**. Expression tracking plots reveal more robust changes in tDR expression (C) when compared with miRNA expression (D) in BeWo cells during $H_2O_2$ treatment;

**E** and **F**. Expression tracking plots reveal more robust changes in tDR expression (E) when compared with miRNA expression (F) in CM cells during $ReO_2$ treatment;

**G** and **H**. Expression tracking plots reveal more dynamic changes in tDR expression (G) when compared with miRNA expression (H) in CF cells during $ReO_2$ treatment.

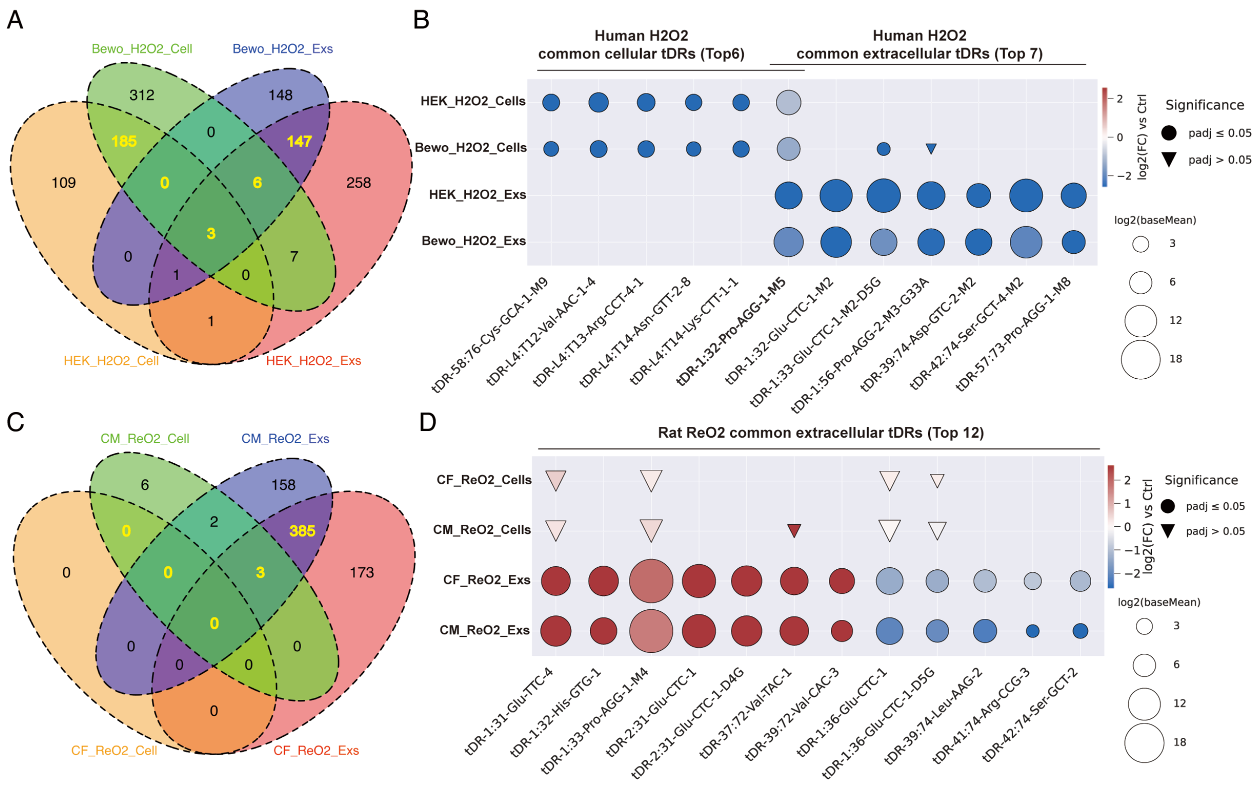


**Figure S14. Oxidative stress-shaped cellular and extracellular tDR signatures**

**A**. Venn diagram shows the overlapped and specific tDRs that were regulated by $H_2O_2$ in human cells and Exs.

**B**. The most significant cellular and extracellular tDRs that were regulated by $H_2O_2$ in both HEK and BeWo cells.

**C**. Venn diagram shows the overlapped and specific tDRs that were regulated by $ReO_2$ in rat cells and Exs.

**D**. The most significant cellular and extracellular tDRs that were regulated by $ReO_2$ in both CF and CM cells.
